# Differences in Alu vs L1-rich chromosome bands underpin architectural reorganization of the inactive-X chromosome and SAHFs

**DOI:** 10.1101/2024.01.09.574742

**Authors:** Lisa L. Hall, Kevin M. Creamer, Meg Byron, Jeanne B. Lawrence

## Abstract

The linear DNA sequence of mammalian chromosomes is organized in large blocks of DNA with similar sequence properties, producing a pattern of dark and light staining bands on mitotic chromosomes. Cytogenetic banding is essentially invariant between people and cell-types and thus may be assumed unrelated to genome regulation. We investigate whether large blocks of Alu-rich R-bands and L1-rich G-bands provide a framework upon which functional genome architecture is built. We examine two models of large-scale chromatin condensation: X-chromosome inactivation and formation of senescence-associated heterochromatin foci (SAHFs). XIST RNA triggers gene silencing but also formation of the condensed Barr Body (BB), thought to reflect cumulative gene silencing. However, we find Alu-rich regions are depleted from the L1-rich BB, supporting it is a dense core but not the entire chromosome. Alu-rich bands are also gene-rich, affirming our earlier findings that genes localize at the outer periphery of the BB. SAHFs similarly form within each territory by coalescence of syntenic L1 regions depleted for highly Alu-rich DNA. Analysis of senescent cell Hi-C data also shows large contiguous blocks of G-band and R-band DNA remodel as a segmental unit. Entire dark-bands gain distal intrachromosomal interactions as L1-rich regions form the SAHF. Most striking is that sharp Alu peaks within R-bands resist these changes in condensation. We further show that Chr19, which is exceptionally Alu rich, fails to form a SAHF. Collective results show regulation of genome architecture corresponding to large blocks of DNA and demonstrate resistance of segments with high Alu to chromosome condensation.

**Graphical Abstract:** 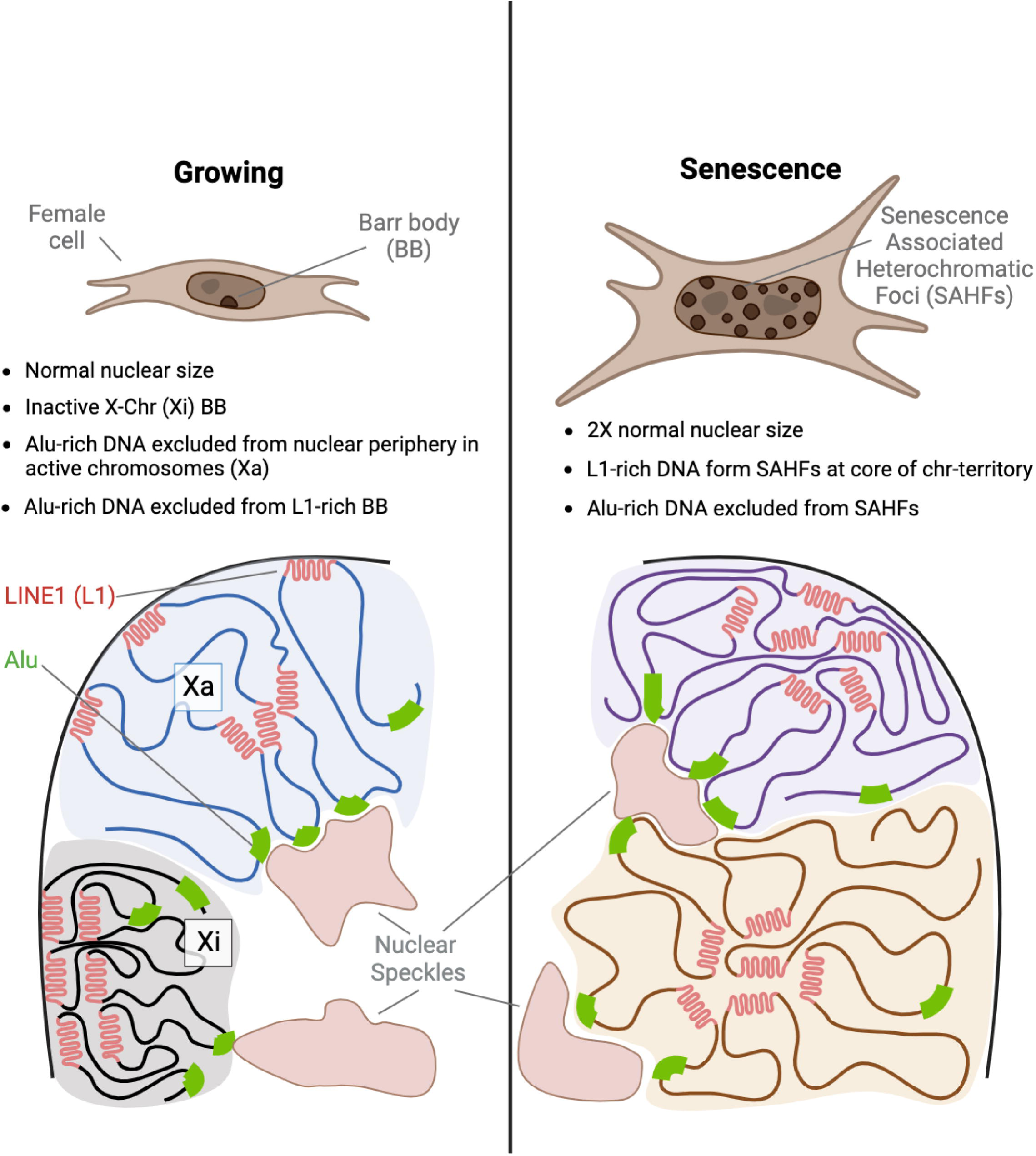

## INTRODUCTION

All cells in the body contain the same DNA yet produce a staggering number of different cell types. This is accomplished by cell-type specific gene expression, controlled through changes in chromatin biochemistry and nuclear structure that impact the *coordinated* activation or silencing of specific gene sets. Much has been learned about the molecular changes involved in regulation of individual genes (e.g. transcription factor binding, nucleosome modifications, etc.), but much less is understood about how the remarkable coordination of ∼20,000 genes in a cell-type specific manner, is achieved. We have long proposed that this higher-level coordination of whole-genome regulation is closely tied to the cytological-scale differences in nuclear organization, with distinct compartmentalization of heterochromatin and euchromatin with other nuclear structures. The nuclear genome organizes in two cytologically distinct chromatin regions; more decondensed euchromatin, within which active genes can express, and condensed heterochromatin, which is primarily silent. These general patterns in spatial organization of the nucleus are cell-type specific, despite some small variations between cells of the same type. As will be discussed below, the large euchromatin compartment is also typically further sub-compartmentalized around ∼20 nuclear bodies that support active gene expression called nuclear speckles.

Despite much progress on understanding nuclear organization at the gene level (Bompadre and Andrey, 2019), the human genome also displays large-scale differences in sequence organization, which are evident cytologically as cytogenetic bands on mitotic chromosomes (discussed below). Importantly, this organization exists even *before nuclear structure forms*, and we suggest may be key to genome function and coordinate regulation. The focus of the work presented here is to investigate whether this large-scale organization of the linear genome is linked to the spatial organization of heterochromatin versus euchromatin, as modeled by whole X-chromosome silencing by XIST RNA, and by genome-wide formation of heterochromatin foci during cell senescence.

Over half of the human genome is made up of highly repetitive interspersed DNA sequences, largely comprised of mobile parasitic DNA elements called SINEs (Short Interspersed Nuclear Elements), and LINEs (Long Interspersed Nuclear Elements). Alu elements are the most abundant SINE in the genome (∼10%), while LINE1 (L1) is the most abundant LINE element (∼17%) (Hoyt et al., 2022). The majority of these mobile repetitive elements in the human genome are degenerate fragments, no longer capable of transposition (reproducing) and are often considered non-functional evolutionary detritus. Interestingly, although the primary sequences of most repeats are rarely conserved across species, their abundance and organization in the genome are conserved.

The organization and variation of sequences across our linear genome can best be seen on mitotic chromosomes which are organized into 2-10 Mb blocks (bands) of DNA that stain differently with certain dyes, most commonly Giemsa-trypsin staining to visualize the “G-dark” and “G-light” band DNA. (Bickmore and Sumner, 1989; Gilbert et al., 2004; Holmquist, 1992; Saitoh and Laemmli, 1994). It has long been known that genes have a clearly non-random distribution relative to these cytogenetic bands, which also differ in GC/AT content, gene density, replication timing and their repeat sequence composition.

For example, “G-dark” bands stain heavily with Giemsa DNA stain, are gene-poor, replicate later in S-phase and are AT and LINE enriched, while “G-light” bands stain weakly with Giemsa, are early replicating, GC, CpG and gene rich, and are enriched for SINE elements. G-light bands are also often referred to as “R-bands” (Reverse bands). (reviewed in: Li and Shen, 2023; Solovei et al., 2016). Routine cytogenetic banding can delineate ∼800 alternating light and dark G-bands, but individual bands may contain smaller segments corresponding to distinct GC or AT isochores (Bernardi, 2021). Why our genome maintains such a large fraction of repeated sequences (∼45%), and why L1 and Alu sequences are organized in cytologically visible mega-structures (chromosome bands) is not well understood.

Each chromosome has a characteristic pattern of these alternating dark and lighter stained bands that is invariant between people and cell-types and has enabled cytogeneticists to use them to identify specific chromosomes and chromosomal abnormalities by microscopy. While chromosome bands have been visualized for clinical purposes for about 60 years, the invariant nature of the cytogenetic banding pattern may have been interpreted to indicate that this high-level structural feature of the human genome is not important for understanding how changes in genome architecture relates to gene regulation. Modern conformation capture analysis uses DNA cross-linking in cells and then extraction of cross-linked DNA to determine spatially proximal DNA, evident on Hi-C interaction maps (Belton et al., 2012). This reveals smaller-scale structural differences in genome organization, evidenced as TADs (topologically associated domains), which are organized within larger clusters in 3D nuclear space, termed A (active) or B (inactive) compartments. Since A & B compartments are generally smaller than metaphase chromosome bands and may show differences between cell-types, there has been little attention to how these smaller levels of organization may relate to large chromosome bands (reviewed in: Solovei et al., 2016).

Interestingly, although sequence conservation between chromosome bands (genes, LINEs or SINES) may not be evolutionarily conserved, repeat composition, Giemsa-staining and syntenic organization of genes within chromosome bands is conserved between species (Bernardi, 2000). Although the functional significance of chromosome banding and its associated repeats is currently unknown, we have long suspected that these blocks of DNA with similar genomic properties may represent a visual manifestation of the DNA framework upon which genome regulation is built, and may explain its evolutionary conservation. We hypothesize the clustering of genes into chromosome bands and their surrounding sequence “fabric” (LINEs, SINEs, GC/AT, etc) facilitates formation of defined nuclear structural compartments, including “closets” of silent heterochromatin or nuclear “hubs” that increase efficiency of gene regulation (Smith et al., 2020).

The mechanisms responsible for this cytological-scale spatial organization of active and inactive chromatin remain largely unknown, although the link between the nuclear lamina and lamina-associated domains (LADs) in heterochromatin have been well studied. However, the role of repeats and the different sequence properties of chromosome bands has in our view been under-studied and often over-looked. In earlier work we examined the nuclear organization of six different chromosome bands and showed that in interphase nuclei, light R-band DNA decondensed and encircled/contacted SC-35 rich nuclear speckles (Shopland et al., 2003). This supported that the clustered gene organization on chromosomes facilitates the interphase organization of active genomic regions in nuclear “neighborhoods” around these non-chromatin hubs (nuclear speckles), rich in pre-mRNA metabolic factors. In contrast, dark G-band DNA was more condensed at the nuclear periphery. In fact, analysis of the chromosomal organization of other types of clustered (tandemly repeated) genes (Chen and Belmont, 2019), supports what we termed the “karyotype to hub hypothesis”, whereby the linear organization of DNA on chromosomes is intimately tied to gene regulation in a highly structured and compartmentalized nucleus (Smith et al., 2020).

Motivated by an interest in how L1-rich dark G-bands and Alu-rich light R-bands relate to changes in genome regulation, we investigate this in two model systems that provide unique synoptic views of how large blocks of L1-rich or Alu-rich DNA rearrange themselves during changes in gene regulation, either across a whole chromosome (X-inactivation), or across the interphase nucleus (cell senescence). X-chromosome inactivation (XCI), the silencing of one X-chromosome in mammalian females, is initiated by a long non-coding RNA called XIST, which is expressed from the inactivating chromosome early in embryogenesis (Brockdorff et al., 1992; Brown et al., 1992). The lncRNA “paints” its parent chromosome (Clemson et al., 1996), and initiates chromosome-wide epigenetic silencing and large-scale structural and nuclear localization changes to the inactivating chromosome. The majority of X-linked chromatin becomes compacted into a densely staining structure called the Barr body (BB), which is then relocated to the heterochromatic compartment at the periphery of the nucleus (or periphery of the nucleolus) in female cells (reviewed in: Brockdorff et al., 2020). Senescent cells also exhibit dramatic rearrangements of euchromatin and heterochromatin across the entire interphase nucleus. When primary somatic cells enter replicative senescence, they irreversibly exit from the cell cycle. And most cell types produce structures called SAHFs (senescence associated heterochromatic foci), where individual chromosomes condense into DAPI bright structures that in many ways resemble the Barr body of the inactive X-chromosome (Narita et al., 2003; Sadaie et al., 2013; Swanson et al., 2013).

## RESULTS

### Relative distributions of Line1 and Alu on chromosomes and nuclei shows marked depletion of Alu-rich DNA in peripheral heterochromatin

We begin this study by examining the overall distribution of L1 and Alu DNA sequences on human mitotic chromosomes and within nuclei of primary human fibroblasts. L1 and Alu sequences are both widely distributed throughout the genome, but are also known to show variations in density in different regions, which correspond generally to cytogenetic bands (Bolzer et al., 2005). As shown in Figure 1A, we used DNA hybridization with probes to Alu (green) and L1 (red), so that the overlay between the two colors highlights differences in their *relative* densities across each chromosome. At this level of resolution, differences in their distributions are apparent as large blocks of DNA (several megabases), that correspond generally to known patterns of chromosome bands (images at right are enhanced)(Fig 1A). Although denaturation and hybridization are known to weaken chromosome banding patterns, you can still see the correlation to Giemsa banding (Fig 1Ai). For example, the characteristic banding pattern of the Chr1 short arm (upper) is a large region of light band DNA, above a region of primarily dark-band DNA: two-color hybridization shows markedly more Alu than L1 in the upper region and more L1 than Alu in the region below that. This difference in sequence organization on the linear DNA of mitotic chromosomes provides perspective for studies below on the relative distributions of L1 and Alu DNA on the inactive X-chromosome (Xi) or other chromosomes within interphase nuclei.

**Figure 1:**
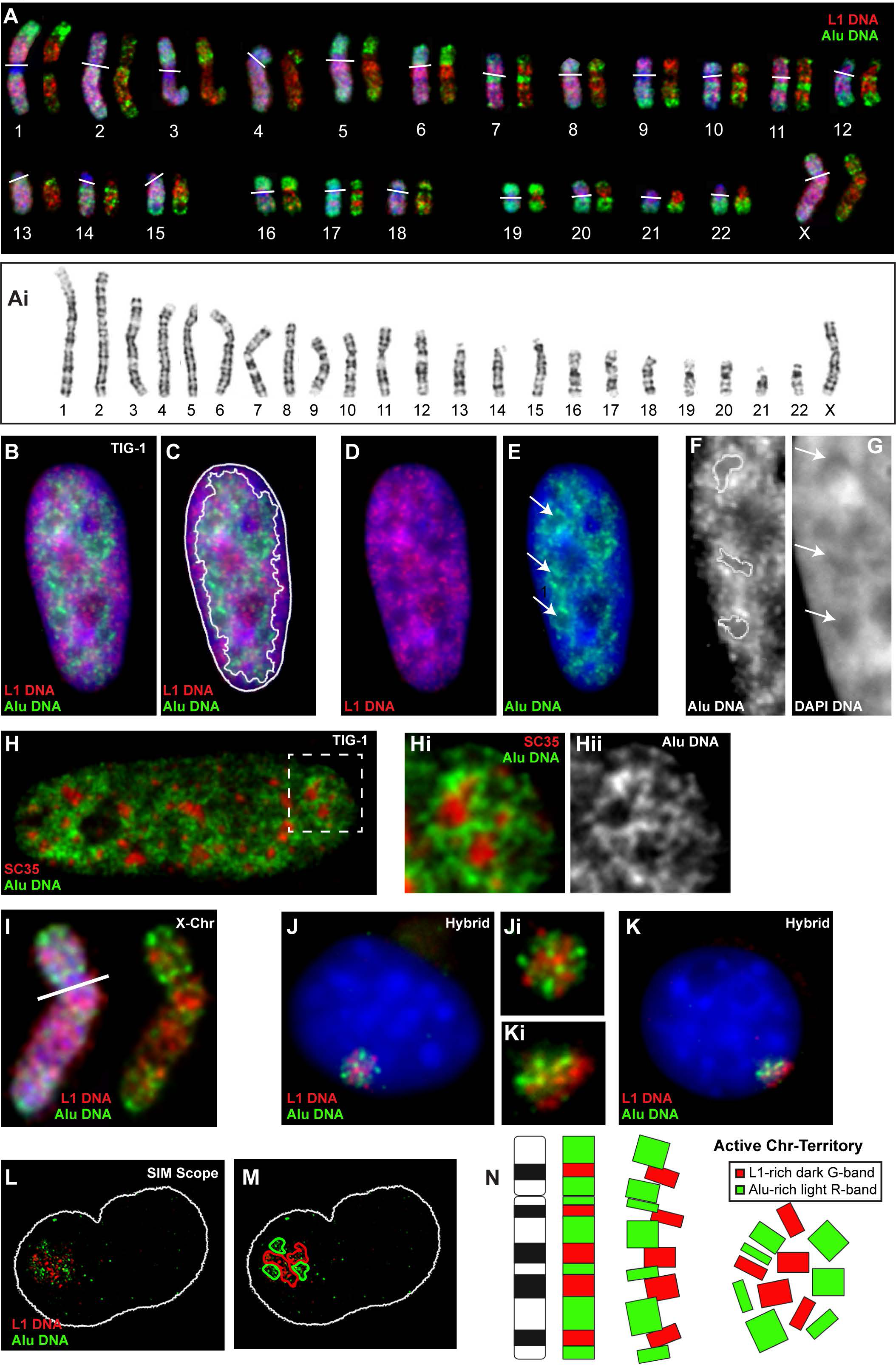
DAPI DNA stain (blue) used in A-G & I-K. **A**) Mitotic human chromosomes with DNA FISH for LINE1 (red) or Alu (green). Left chromosome image are three color channels including DAPI (Blue). Right chromosome images show only two color channels, with enhanced contrast and brightness. White lines at centromeres. Below: standard 550 band human karyotype with Giemsa DNA stain (**Ai**). **B-G**) Human fibroblast nucleus with DNA FISH for LINE1 (red) or Alu (green). Area of green signal outlined in (**C**). Region indicated by arrows in (**E**), is enlarged in (**F&G**). Alu-DNA “rings” and DAPI “holes” are indicated by arrows (**E&G**) or outlined in (**F**). **H**) Human fibroblast nucleus with DNA FISH for Alu (green) and IF for SC35 protein (nuclear speckles) (red), with square region enlarged in (**Hi**). **Hii**) is green channel of enlarged region (**Hi**). **I**) Human X-chromosome from Figure 1A, line is at centromere. **J-K**) Mouse/human hybrid cells containing single human X-chromosome with DNA FISH for human LINE1 (red) or Alu (green). Close-up of chromosomes in small images at center. **L-M**) Super-resolution structured illumination microscope (SIM) image of mouse/human hybrid X-Chr cells with DNA FISH for human LINE1 (red) or Alu (green). Regions of mostly red or green signal outlined at right. **N**) Diagram illustrating the potential clustering of chromosome band regions enriched for L1 (red) or Alu (green) in an active interphase chromosome territory.

L1-rich DNA is reported to be enriched at the nuclear periphery of interphase nuclei due to the density of lamina-associated domains (LADs), with some published images showing much or most L1 DNA at the nuclear periphery with little L1 in the internal part of the nucleus (e.g. Lu et al., 2021; Solovei et al., 2016); however the extent to which this is seen will vary with cell-type and with digital imaging and processing procedures. We find the enrichment of L1 DNA at the nuclear periphery of human fibroblasts, is not particularly striking (e.g. Figs 3C-D), and even in cells with a well-defined peripheral L1 compartment, (Fig 1B-E), it actually appears to be determined more by the *exclusion of Alu DNA* than increased L1. For example, in images where L1 looks significantly enriched at the nuclear lamina in three-color images (Fig 1B-C), L1 does not look to be particularly increased at the periphery in single-channel images (Fig 1D). On the other hand, the signal for Alu DNA is largely *excluded* from that peripheral region (Fig 1E), with minimal photographic manipulation (see methods). This was seen in these normal primary cells but was also apparent to a lesser extent in other cell lines examined below (Fig 3E). The depletion of Alu at the nuclear periphery would not be inconsistent with studies associating L1 sequences with LADs, but does introduce a conceptually distinct possibility: that lack of Alu, rather than enrichment of L1 sequences, may be a hallmark of the peripheral heterochromatin compartment, in normal human fibroblasts and potentially other cells.

The marked depletion of Alu-rich DNA from the peripheral heterochromatin suggests Alu exhibits a more specific relationship with nuclear compartments than does L1 DNA. In fact, within the interior euchromatic regions we find brightly staining rings of Alu DNA surrounding area depleted of DNA (Fig 1E arrows & Fig 1F-G). These areas, which appear as dark holes in the DAPI stain, contain nuclear speckles (SC35 domains) (Fig 1H), which are non-chromatin, membraneless nuclear structures, highly enriched for poly-A RNA and pre-mRNA splicing factors. We previously showed that many active genes, including very highly expressed genes, specifically localize and are transcribed at the outer edge of an individual speckle, and various post-transcriptional processes can occur within each of these “factories” (Smith et al., 1999; reviewed in: Hall et al., 2006). More recent studies affirm the preferential distribution of most active genes with speckles occurs genome-wide (Chen and Belmont, 2019). We have previously suggested that the preferential localization of protein-coding genes in light R-band (Alu-rich) DNA may facilitate this clustering of genes with nuclear hubs that facilitate their expression (Smith et al., 2020) and showed that the three light R-bands examined associate with nuclear speckles, whereas the three dark G-bands did not (Shopland et al., 2003). Supporting this hypothesis more broadly, here we show that Alu-rich DNA across the genome, a loose proxy for R-band DNA, is preferentially associated with nuclear speckles (Fig 1H).

To gain insight into L1 and Alu distributions within an individual interphase chromosome territory, we briefly examined mouse/human hybrid cells that contain a single active human X-chromosome (Xa) (Fig 1I). Since primary sequences of repeats are poorly conserved, *human* L1 and Alu FISH probes are specific to the human chromosome. In these hybrid nuclei, we observe L1 and Alu DNA clusters that appear largely segregated within the human chromosome territory (labeling red or green but not yellow) (Fig 1J-N). In most (∼90%) cells the clusters of L1- and Alu-rich DNA exhibit a random distribution across the territory (e.g. Fig 1J), while in a smaller fraction (10%) in which the chromosome appeared compressed against the nuclear periphery, Alu DNA was more towards the interior and L1 against the nuclear envelope (n=55-100 in three experiments)(Fig 1K).

Thus, at the intrachromosomal level several distinct clusters of more L1-rich or Alu-rich DNA signal are seen within an individual territory, which may largely reflect their chromosomal distribution and/or the homotypic clustering that has been suggested based on Hi-C (Lu et al., 2021)(Fig 1N). However, at the level of the overall nucleus in normal human fibroblasts we find broader distribution of L1 DNA than Alu DNA and suggest that a more defining feature of peripheral heterochromatin may be lack of Alu, shifting the relative densities of Alu to L1. The preferential distribution of Alu-rich DNA in the interior is associated with, and may be driven by, the close association of gene-rich DNA with nuclear speckles, which distribute in a central plane within the interior of fibroblast nuclei (Carter et al., 1993).

To further investigate the organization of Alu-rich R-band versus L1-rich G-band DNA, we examined this in two natural systems in which whole chromosomes undergo dramatic changes in architecture and regulation.

### Alu/gene-rich and L1-rich DNA are organized differently in chromosomes silenced and condensed by XIST RNA

When the lncRNA XIST “paints” and silences the X-chromosome in female cells, it forms a large heterochromatic structure called the Barr body (BB), that in human cells is visible in most cells by DNA stains like DAPI (Fig 2A arrow). This DAPI-dense BB is often presumed to comprise the entire X-chromosome, compacted by the silencing and condensation of almost all genes across the chromosome. We previously showed that hybridization with a probe to CoT-1 DNA (the highly repetitive fraction of the genome) brightly marks the BB (Clemson et al., 2006), which is devoid of CoT-1 RNA (Fig 2B). However, we were surprised to find that 14 individual genes, regardless of activity (silenced or escaped), did not distribute within the DAPI-dense BB structure, rather the coding genes primarily localized at the outer periphery. Nonetheless, it is still often presumed that the BB represents the whole inactive chromosome containing all the silenced genes. Therefore, we briefly repeated gene localizations that illustrate and affirm the prior more quantitative and extensive analysis. Figure 2C-I shows representative localizations for five different genes. In fact, DNA FISH using a pool of four active and inactive X-linked gene probes shows the collection of genes encircling the very outer edge of the XIST RNA cloud over the BB (Fig 2G-I). Clemson *et al* found the BB forms an inner core of the inactive chromosome territory, but is smaller than the XIST RNA cloud and the Xi territory seen with a DNA paint (Clemson et al., 2006; reviewed in: Hall and Lawrence, 2010). Thus, during the silencing process, XIST RNA reorganizes the inactive chromosome territory such that the center core of the territory is composed of a highly condensed ball of repeat-DNA (the Barr body) surrounded by mostly silent and some active genes.

**Figure 2.**
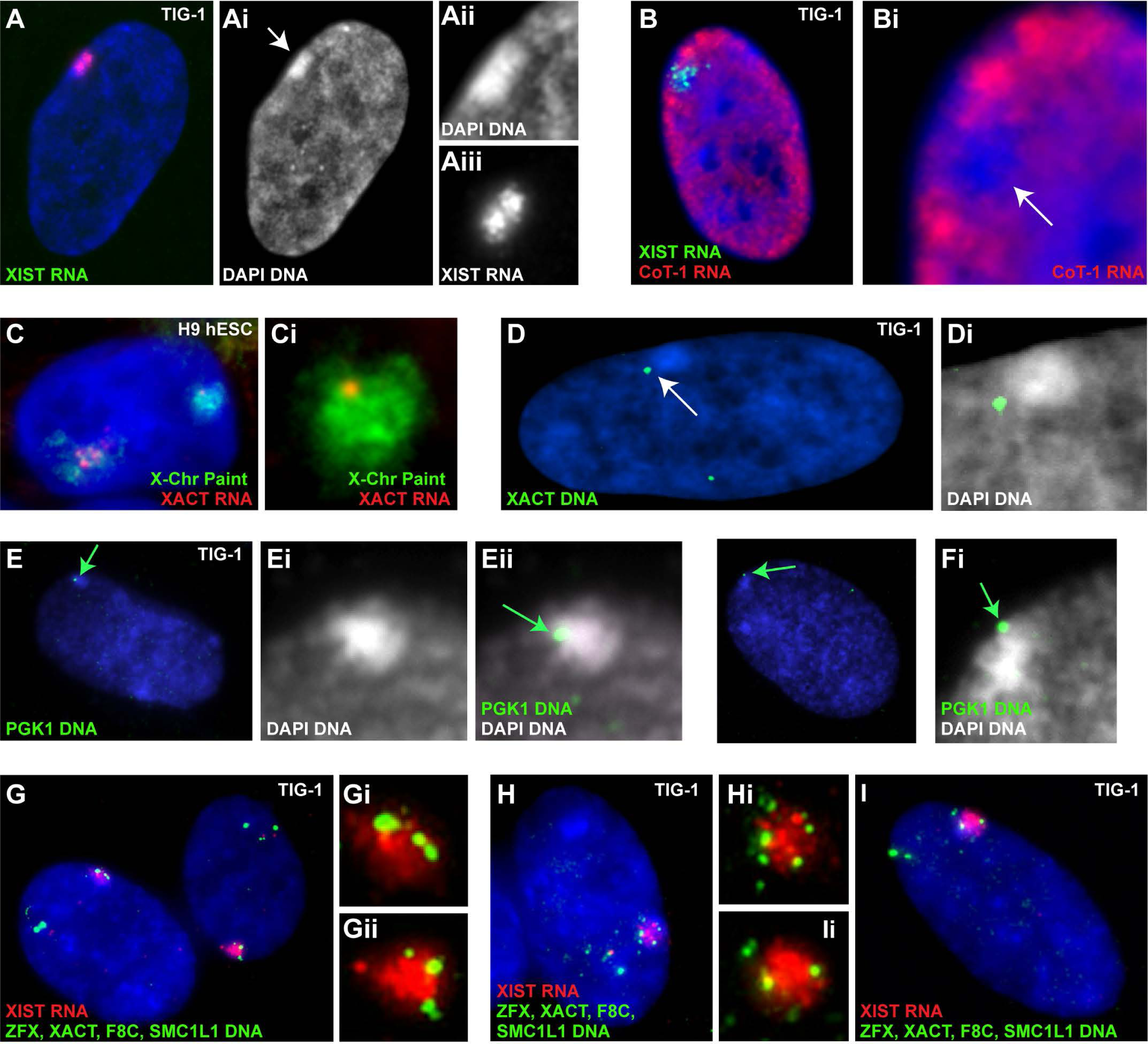
DAPI DNA stain (blue) used in all images. **A**) XIST RNA FISH (red) is coincident with DAPI dense Barr body (BB) (arrow) in female fibroblast. Single channel close-ups of region at right. **B**) Female fibroblast with XIST RNA (green) and CoT-1 (red) hnRNA FISH. CoT-1 RNA “hole” (arrow) over Xi enlarged at right, without XIST RNA signal. **C**) Human H9 ES cell with XACT RNA FISH (red), and X-chromosome paint DNA FISH (green), and two-color close-up of Xi in insert at right. **D**) Female fibroblast nucleus with XACT DNA FISH (green). Enlarged image of BB with DAPI DNA (white) and XACT gene signal (green) at right. **E-F**) Female fibroblast nucleus with PGK1 DNA FISH (green) and enlarged regions of BB (DAPI: white) with gene (green) indicated by arrow at right. **G-I**) Female fibroblasts with a pool of four X-linked genes labeled by DNA FISH (green) and XIST RNA FISH (red). Two-color close-ups of Xi next to all images. X-linked gene activity state: **PGK1** = silenced, **ZFX** = escape, **XACT** = silenced, **F8C** = silenced, **SMC1L1** = silenced.

Enrichment of the CoT-1 DNA probe in the BB of female cells suggests high-copy repeats are abundant in this structure. And as expected, we find the human BB labels brightly with L1 probes (L1 enrichment in 94% of BBs: stdev 5)(Fig 3A-B). In fact, the labeling intensity of L1 DNA over the BB is striking, making the BB stand out as generally the biggest and brightest L1-rich domain in the nucleoplasm. This suggests, the BB contains a significantly higher L1 DNA density (or organization) than any other DNA in the nucleus including L1-rich DNA in the peripheral heterochromatic compartment.

**Figure 3.**
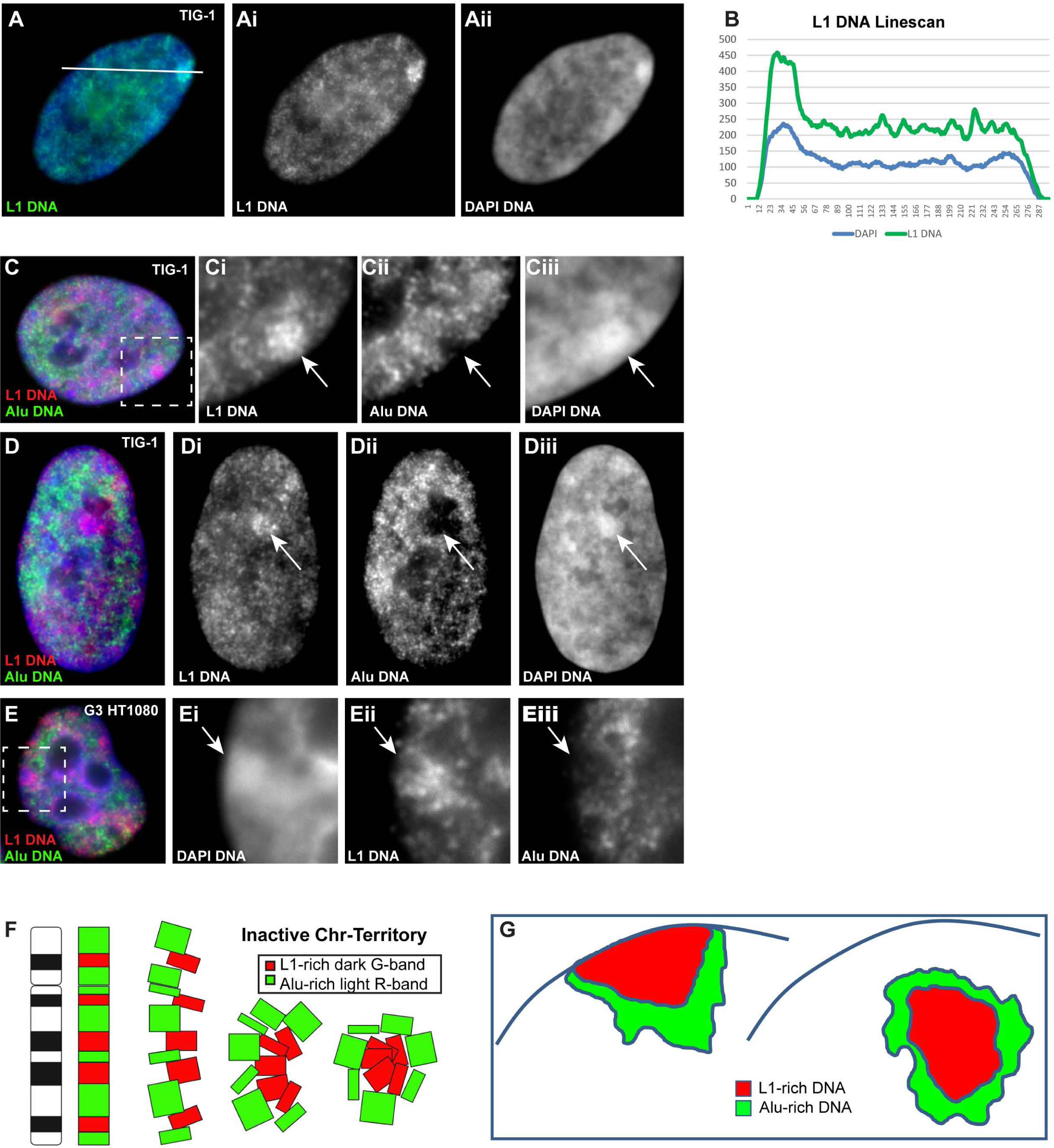
DAPI DNA stain (blue) used in all images. **A-B**) Female fibroblast with L1 DNA FISH (green). Single-color channels separated at right. **B**) Linescan histogram of green and blue intensity levels for line shown in (**A**). **C-D**) Two examples of female fibroblasts with L1 DNA (red) and Alu DNA (green) FISH. At right, separated channels of region indicated by square (C) or whole image (D), with Xi indicated by arrow in each channel. **E**) Transgenic G3 HT1080 cell with L1 DNA (red) and Alu DNA (green) FISH. At right, enlarged and separated channels of region indicated by white box, with inactive chromosome indicated by arrow in each channel. **F**) Diagram illustrating the clustering of L1-rich chromosome bands (red) into the center of the inactive chromosome territory, surrounded by Alu-rich (green) band regions. **G**) Diagram illustrating the two domains of the inactive chromosome, enriched for L1 (red) or Alu (green), and localized against the nuclear membrane or nuclear interior.

Although L1 specific probes have often been used as a proxy for CoT-1 DNA to label the BB in the past (e.g. Chow et al., 2010), to our knowledge whether Alu-rich DNA is comparably condensed into the BB has not been examined, hence it’s possible that not all high-copy repeats, and chromosomal regions, participate equally in BB formation. Using DNA FISH, we examine whether Alu-labeled DNA is enriched within the repeat-rich BB, along with L1 DNA. Remarkably, we find that Alu DNA was not enriched in the BB in 100% of nuclei scored (e.g. SuppFig1A), and very often there was a region from which Alu signal was largely *excluded* (79%: Stdev 16), appearing as a “hole” in the nucleoplasmic Alu DNA signal over the BB (Fig 3C-D: arrows). The difference in L1 and Alu signals was not due to difficulty in denaturing the condensed BB, as the L1 DNA clearly hybridized, nor was it due to differential probe penetration as both probes were nick-translated into similar fragment size. Instead, results suggest that chromosome condensation of the L1 dense BB structure triggered by XIST RNA, nucleates in a region substantially depleted of Alu sequences. This is similar to Alu exclusion in the nuclear periphery (Fig 1B-E), but occurs irrespective of whether the BB is associated with the nuclear envelope (Fig 3D), suggesting LADs/Lamina association does not play a role. And, because Alu sequences are enriched in gene-rich R-bands, this also supports our findings that genes are located outside the BB structure (Clemson et al., 2006) (Fig 2C-I). This differential organization of L1 and Alu sequences was also seen in the BBs of autosomes silenced by XIST RNA (e.g. Chr-4 in G3/Ht1080 sarcoma cells: Fig 3E) (Hall et al., 2002). This further supports our previous findings that autosomes exhibit the same pattern of genes located outside the BB (Clemson et al., 2006) and suggests that this L1/Alu DNA organization (Fig 3F-G) may be a universal structure generated by XIST-mediated silencing.

Results demonstrated above indicate that within a silenced chromosome territory there is congregation of the homotypic intra-chromosomal DNA clusters marked by Alu versus L1-rich DNA. Figure 1 shows the banding patterns of L1 and Alu DNAs on human chromosomes, and Figure 3F graphically illustrates these DNA blocks with high L1 and low Alu congregating to form the condensed Barr body. Further supportive of this, in a mouse human hybrid cell carrying a human inactive X-chromosome (Xi) (Brown and Willard, 1994), the several smaller clusters of L1 versus Alu rich DNA that were seen for the active X in a hybrid cell (Fig 1J&N) are now congregated into two distinct L1 and Alu-rich domains (SuppFig1B-D). Thus, chromosome silencing by XIST RNA induces a reorganization of L1 versus Alu-rich DNA into two regions within the chromosome, and a similar segregated organization is largely retained even in hybrid nuclei of a different species.

Interestingly, we note certain cytological observations which suggest that XIST RNA may have a differential affinity for the L1-rich and Alu-rich regions of the Xi. For example, as Xist RNA falls off mitotic chromosomes in mouse cells, it can be seen painting the *entire chromosome* in very early prophase, including L1-rich G-bands, (SuppFig 2A: (Smith et al., 2004)) but is retained longest on Alu-rich R-bands later in metaphase (SuppFig 2B). Similarly, when *XIST* transcription is compromised (e.g. transcriptional inhibitors), XIST RNA is retained longest on the DNA around the edge of the BB where we find Alu-rich DNA resides (SuppFig 2C-F) (Clemson et al., 1996). Thus, under certain conditions XIST RNA may show differences in binding affinity or stability relative to gene-poor/L1-rich DNA versus gene-rich/Alu-rich DNA. However, at both interphase and early mitosis the RNA is seen to coat essentially the whole chromosome, so differences in binding/retention should not be conflated with its normal association with the chromosome territory in interphase.

As discussed in the supplement, we also examined XIST RNA labeling relative to H3K27me3 and H3K9me3 heterochromatin marks, given some earlier reports that the Xi is segregated into two domains differentially marked by H3K27me3, H3K9me3 and XIST RNA (Chadwick and Willard, 2004; Nozawa et al., 2013). These reports suggested that XIST RNA does *not* bind the entire Xi territory and avoids H3K9me3 labeled chromatin (Dixon-McDougall and Brown, 2016), which is known to mark L1-rich DNA. However, here we find that XIST RNA is clearly and consistently associated with the DAPI-dense BB, which is enriched for H3K9me3 (SuppFig 2G-H), H3K27me3 (SuppFig 2I), and L1 DNA (SuppFig 2J). Importantly, we found that choice of XIST probe can compromise antibody labeling. H3K9me3 labeling followed by XIST RNA hybridization using a 9kb 3’ XIST probe (G1A), or vice versa, compromises the second signal (SuppFig 2K), replicating reports of mutually exclusive XIST RNA and H3K9me3 labeling. However, when a full-length XIST cDNA probe (∼15bk) is used, more complete overlap between the two signals is seen (SuppFig 2H). Thus, reports suggesting H3K9me3 and XIST RNA are mutually exclusive might be impacted by XIST RNA retention, probe choice, labeling method and antibody used, and we believe XIST RNA does paint the L1-rich, H3K9me labeled, BB DNA as well as silenced Alu/gene-rich DNA.

Thus, the BB is not the whole condensed X-chromosome with all repeat-rich DNA similarly represented. Alu-rich DNA is significantly excluded from the BB, which supports the finding that most genes regardless of activation state are located outside the structure of the BB. And, XIST RNA paints all silenced Xi chromatin, although it might exhibit different affinity or stability on the two regions.

### SAHFs are similar to the BB and further suggest that relative L1/Alu density impacts chromosome compaction

We next examined the respective distributions of Alu and L1 DNA in relation to large-scale changes in heterochromatin organization that occurs genome-wide in cell senescence. The nuclei of senescent cells enlarge by about two-fold, their centromeric DNA unravel and string-out (Swanson et al., 2013), and in most cell types the chromatin forms numerous DAPI dense structures called Senescence Associated Heterochromatin Foci (SAHFs)(Fig 4A-C). As previously shown, SAHFs are marked by a lack of the heteronuclear CoT-1 (repeat-rich) RNA which brightly labels the rest of the decondensed, euchromatic nucleoplasm (Creamer et al., 2021). Hence, SAHFs are similar to the BB in that they comprise silenced heterochromatin that lacks RNA expression (e.g CoT-1 RNA detects repeats in pre-mRNA introns and lncRNAs) (Fig 4D-E).

**Figure 4:**
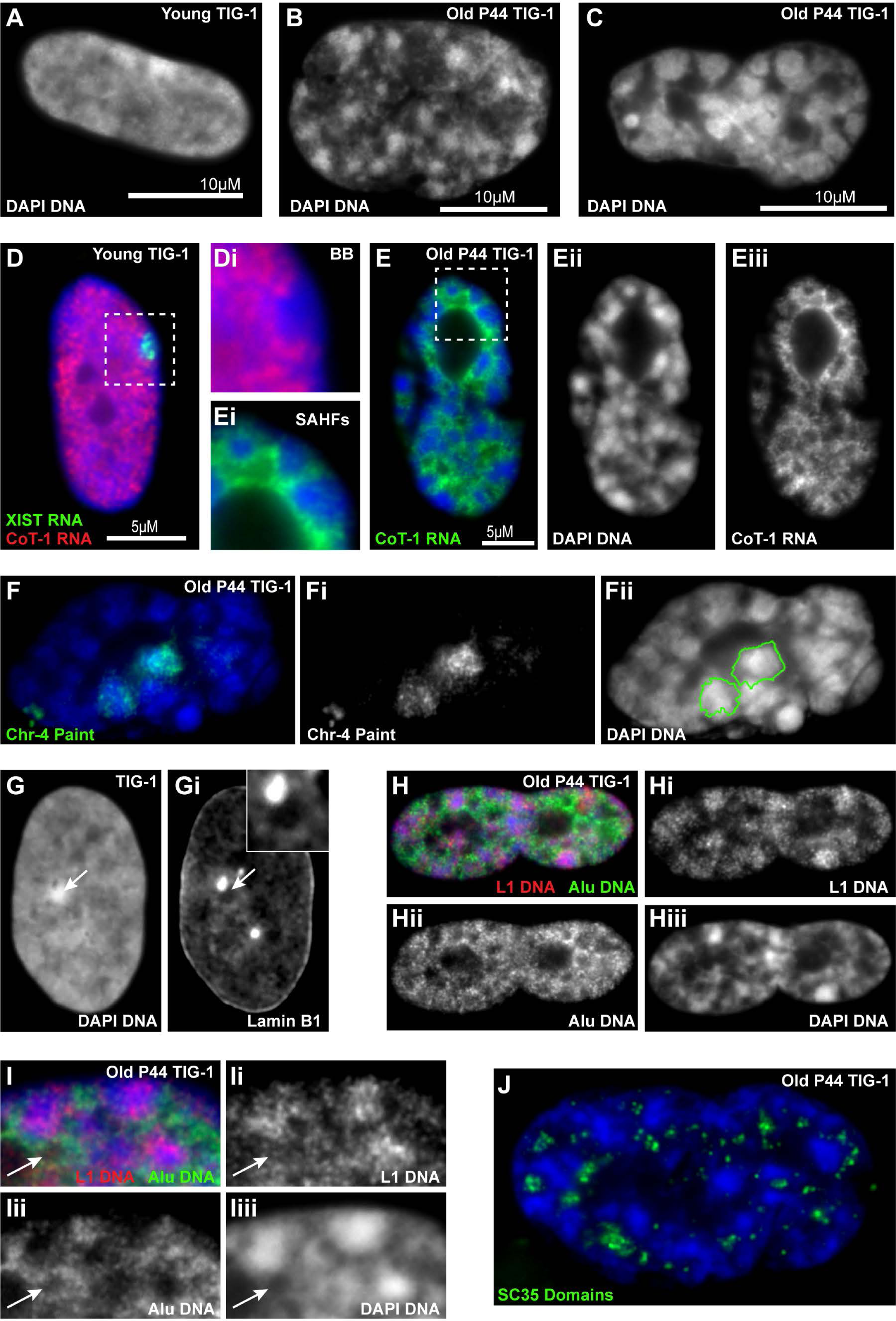
DAPI DNA stain (blue) used in all images. **A-C**) Single channel DAPI images of growing (**A**) or senescent (**B-C**) fibroblasts, with varying sizes of DAPI dense SAHFs in senescent cells. **D**) Female fibroblast with CoT-1 (red) and XIST (green) RNA FISH. Enlarged region indicated by white box (at right), without XIST RNA to show CoT-1 RNA “hole” over BB. **E**) Senescent fibroblast with CoT-1 RNA (green). Separated channels at right and enlarged region indicated by white box (at left), to show CoT-1 RNA “holes” over SAHFs. **F**) Senescent fibroblast with Chr-4 DNA FISH (green), with separated channels at right and outlines of both Chr-4 territories superimposed on DAPI image. **G**) Separated channels of DAPI DNA stain (left) and Lamin B1 staining (right) in a female fibroblast, with BB indicated by arrow. Insert is an enlargement of region over BB, from Lamin B1 channel. **H**) Senescent fibroblast with L1 (red) and Alu (green) DNA FISH. Separated channels included. **I**) Close-up of SAHFs from a senescent fibroblast with L1 (red) and Alu (green) DNA FISH and separated channels. Arrows indicate Alu-DNA “ring” around DAPI “hole”. **J**) Senescent fibroblast with SC35 protein (nuclear speckles) labeled in green.

Given this, we examined whether transcriptionally silent, DAPI-bright heterochromatic SAHFs also represent the inner “core” of a single chromosome territory like the BB. We examined this for three chromosomes using whole chromosome DNA paints as probes in replicatively-senescent human fibroblasts (P44), with ∼50 or more cells examined for each. Representative images are shown to illustrate the consistent observation that in most cells a single SAHF occupied the center of a single chromosome territory, even for very large SAHFs (e.g. Fig 4F & SuppFig 3). This is consistent with a suggestion from two earlier studies (Funayama et al., 2006; Zhang et al., 2007). Hence, most SAHFs represent a condensed core of the chromosome territory largely analogous to the Barr Body. Importantly, the SAHF does not encompass the entire chromosome territory; since SAHFs are surrounded by especially decondensed DNA in the enlarged nucleus, some less condensed territory DNA can be seen outside the condensed structure (Fig 4F: green outline, and SuppFig 3), similar to the BB (reviewed in: Hall and Lawrence, 2010).

It is known that in senescent cells there is a loss of Lamin B1 associated with rearrangement of peripheral heterochromatin to form SAHFs (Sadaie et al., 2013; reviewed in: Lukasova et al., 2018). In non-senescent cells Lamins are not only concentrated at the periphery but are also present across the nucleoplasm. Interestingly, in cycling cells with normal Lamin levels we noted diminished nucleoplasmic Lamin B1 staining associated with the BB (Fig 4G and SuppFig 4), suggesting another parallel between formation of these two chromosomal “core” structures. Loss of CoT-1 repeat RNAs from euchromatin (as triggered by XIST RNA) causes chromatin condensation associated with disruption of an insoluble RNA-protein scaffold (Creamer et al., 2021). Hence, depletion of Lamin B1 association may be a common feature of both BB formation as well as SAHFs.

These similarities between BBs and SAHFs suggest that SAHFs might also exhibit similar L1/Alu compartmentalization as the BB. Using L1 and Alu DNA FISH on SAHF-containing senescent fibroblasts, we find the same L1-enrichment and Alu-depletion over DAPI-bright SAHFs; the consistency of this pattern is evident for SAHFs throughout a nucleus (Fig 4H). These results support that the intrachromosomal congregation of chromosome segments with similar L1/Alu content, which form a bipartite segregated structure, is not exclusive to XIST-mediated chromosome silencing. The Alu-depleted peripheral heterochromatic compartment depleted of Alu seen in cycling fibroblasts (Fig 1B-C) is less apparent in senescent fibroblast nuclei. Instead, the L1-rich DNA that lacks substantial Alu appears to have collapsed into the center of each chromosome territory to form a SAHF. In contrast, Alu-rich regions remain largely outside these L1 dense structures, and Alu-rich DNA closely associates with non-chromatin speckles rich in SC35 and other RNA metabolic factors (Fig 4H-J: arrows).

Because DAPI-bright SAHFs and BBs are formed from DNA low in Alu and rich in L1, this raised the possibility that chromosomes with higher relative L1/Alu density might have enhanced SAHF or BB formation. Therefore, we considered the overall content of L1 versus Alu for all human chromosomes, as graphed in Figure 5A. The human X-chromosome (except the region that escapes silencing) is highly L1-rich, and this may contribute to its capacity for silencing by XIST RNA (reviewed in: Lyon, 2006). Chr-4 and Chr-19 are standout autosomes with the highest L1 and Alu concentrations respectively, as also evident by DNA FISH (Inserts Fig 5A), hence comparison of those two chromosomes for SAHF formation was of interest. Related to this, we note that our human cell line carrying an XIST transgene on L1-rich Chr-4 (G3/HT1080)(Hall et al., 2002) produces a particularly large, bright and round BB (Fig 5B). SAHF formation for Chr-4 and Chr-19 in senescent fibroblasts was determined by size and intensity of DAPI stain and approximate percent of chromosome territory occupied by the SAHF. To define chromosome territories, we used labeled probe libraries to Chr-4 and Chr-19, which are optimized for cytogenetic chromosomes preparations and not our standard fixation of interphase nuclei. Thus, the highly decondensed outer edges of labeled chromosome territories may be harder to visualize, and all measurements underestimate full territory size to some degree. We find 100% of Chr-4 territories make a large SAHF in senescent nuclei, and it was usually one of the largest and brightest SAHFs (Fig 5C: arrows, 5F and SuppFig 3A-B)(70% of the biggest or brightest SAHFs were Chr-4), second only to the Xi. In addition, the Chr4-SAHF encompassed the majority of the Chr-4 paint territory (Fig 5C: outline & 5G). Chr-19, on the other hand, localizes in DAPI-dim decondensed nuclear regions in growing cells (Fig 5D: outline), and in senescent cells only about 50% of the Chr-19 territories contained even a tiny DAPI dense region that could represent a small SAHF (Fig 5E: outline). The tiny DAPI-regions of Chr-19 were very dim and occupied only about 20% of the chromosome territory (Fig 5G). Thus, there are marked differences in the frequency and extent of SAHF formation, suggesting that the L1 and Alu density of different chromosomes may play a role in SAHF/BB formation.

**Figure 5:**
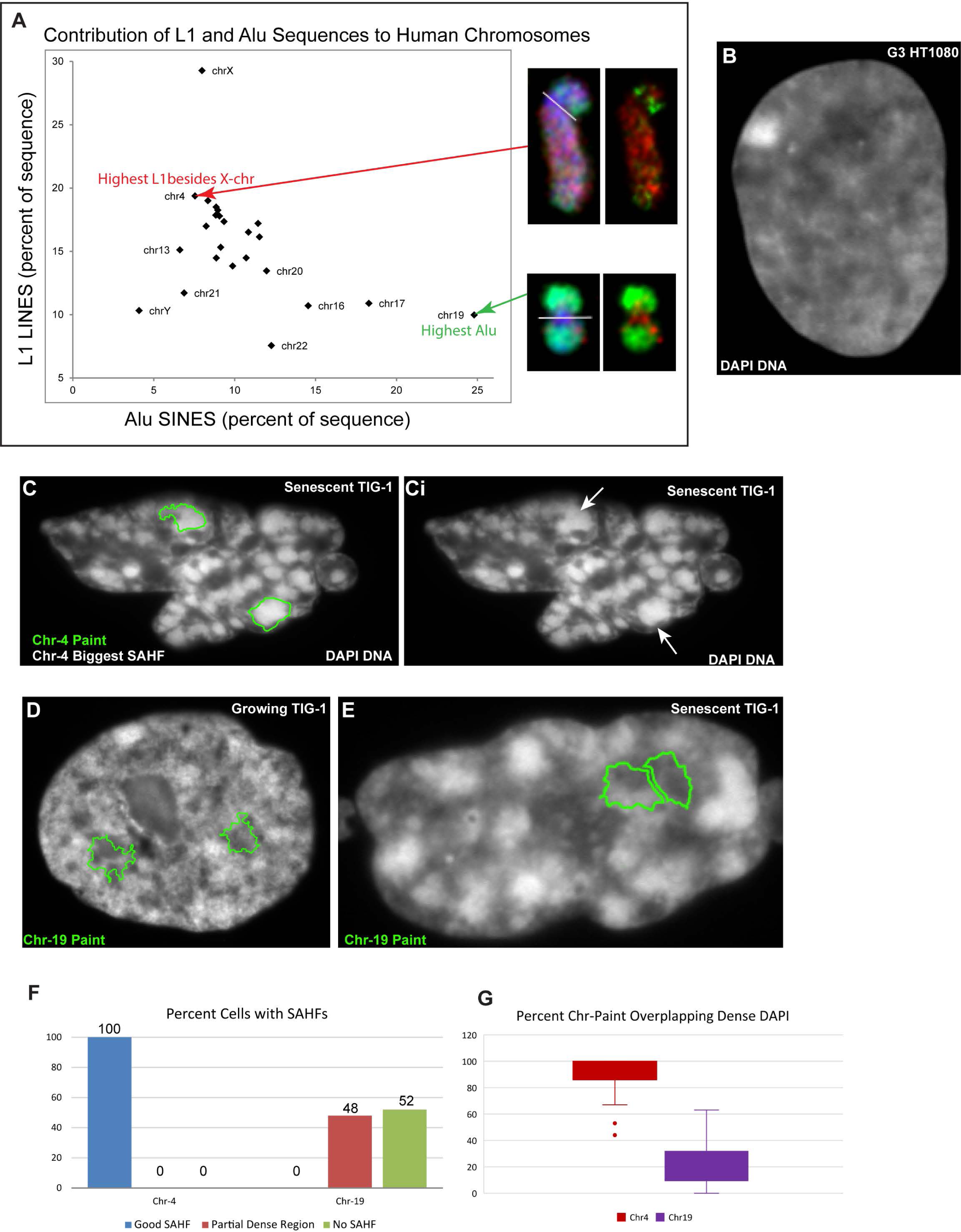
**A**) Graph of L1 and Alu sequence contribution (percent of total chromosome sequence) for all human chromosomes. Inserts of Chr4 and Chr19 with L1 (red) and Alu (green) DNA FISH from Figure 1A. **B**) DAPI DNA stain of Transgenic G3 HT1080 nucleus with XIST integrated into Chr4. **C**) DAPI DNA stain of senescent fibroblast nucleus with outline of both Chr4 DNA paint territories superimposed on DAPI image on left, and arrows indicating the same two SAHFs at right. **D-E**) DAPI DNA stain of young (**D**) and senescent (**E**) fibroblast nuclei with outline of both Chr19 DNA paint territories superimposed on DAPI images. **F**) The presence, size and intensity of SAHFs in senescent fibroblasts were scored for 100 cells per Chr-paint experiment. **G**) The areas of SAHF and Chr-paint territories were compared and measured for the percentage of Chr-paint that overlapped the SAHF.

We conclude that DAPI-dense SAHFs in senescent cells share many characteristics with the inactive BB, including lack of significant CoT-1 RNA transcription, high L1-DNA density, the exclusion of Alu-rich DNA, and decreased Lamin B1 association. In addition, with parallels to the BB, SAHFs do not represent an entire compacted chromosome territory, but constitute the central heterochromatic L1-rich core of a single chromosome territory.

### Hi-C reveals L1-rich DNA gains distal *intra*chromosomal interactions as G-bands compact into SAHFs but Alu-rich segments resist

As conceptualized in Fig 6A, the above cytological results indicate that in senescent cells the architecture of individual chromosomes changes dramatically as distal but syntenic L1-rich/Alu-poor regions coalesce together to form a SAHF in the center of the territory. At the same time, results suggest that highly Alu-rich regions would not coalesce into the SAHF but remain more distended in the peripheral regions of the territory, in a much larger senescent nucleus. To examine the changes on individual chromosomes at higher sequence resolution, we began by examining changes in Hi-C intrachromosomal interactions between growing and senescent human cells using published data (Chandra et al., 2015). (We note that Chandra et al., 2015 examined *inter-* chromosomal long-range interactions broadly across the genome, but not specifically *intra*-chromosomal, see Discussion.)

**Figure 6:**
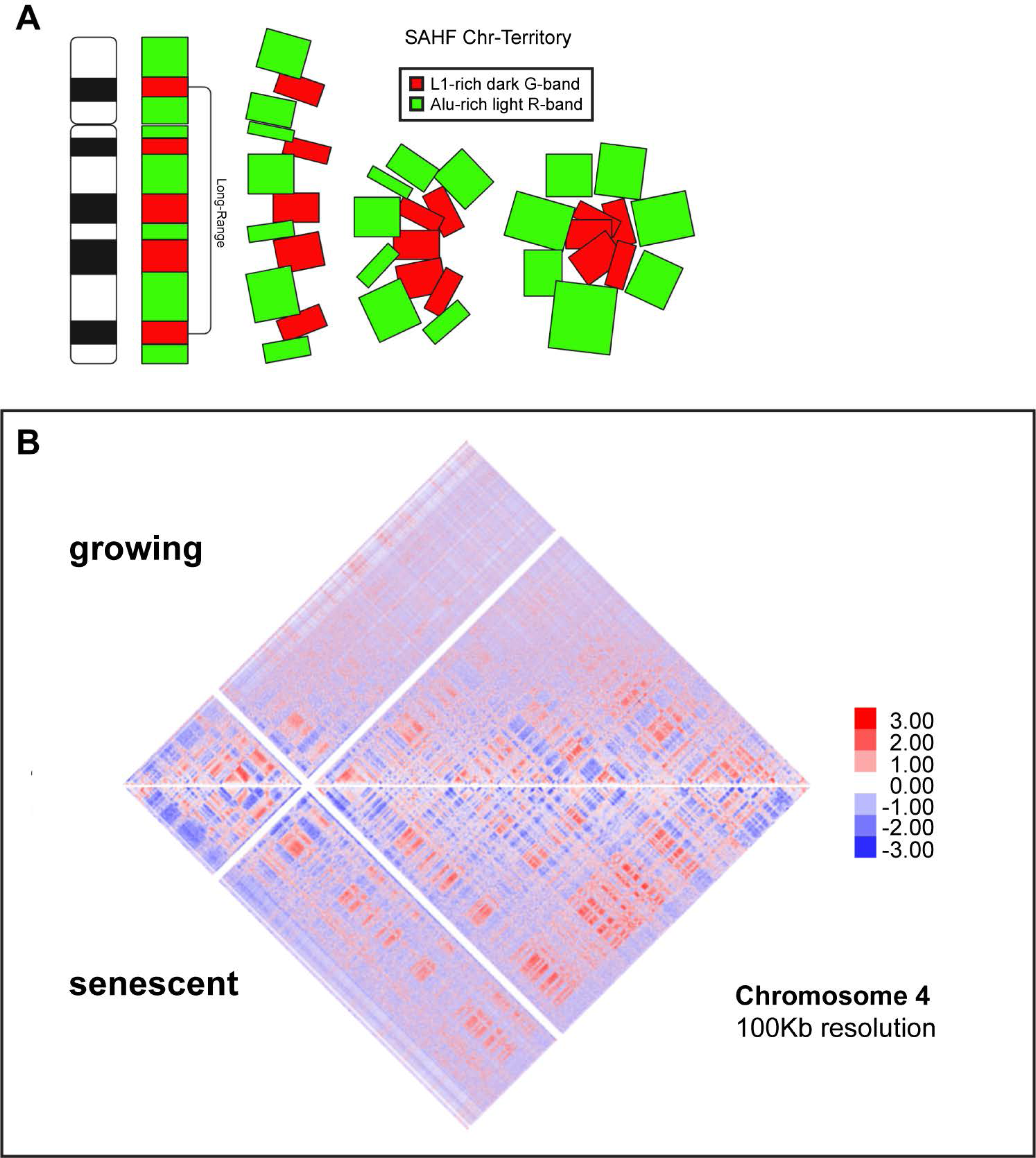
**A**) Diagram illustrating the clustering of chromosome band regions enriched for L1 (red) condensed into the center of the senescent chromosome territory SAHF, picking up syntenic long-range interactions, and surrounded by distended Alu (green) band regions. **B**) Hi-C interaction heatmap of Chromosome 4 in growing (top) and senescent (bottom) fibroblasts at 100kb resolution (red = gains, blue = losses).

Our analysis and findings bear on three inter-related questions/hypotheses. First, we hypothesized that L1-rich regions would pick up new long-range *intrachromosomal* interactions, thus increasing the proportion of interactions that are long-range (>10Mb) versus short-range (>10 Mb). Second, we examined the extent to which large contiguous blocks of DNA (several megabases or more) would change in concert, particularly in regions of light versus dark cytogenetic bands. Third, we examine whether L1 and/or Alu density correlates with and may influence architectural remodeling, and in particular examine the top 10% L1 rich versus top 10% Alu rich DNA.

Figure 6B shows interaction maps for growing versus senescent cell DNA for human Chr-4 (blue are losses and red are gains), which above results show forms a very prominent SAHF. Increased long-range syntenic interactions are evidenced by the prevalence of red patches (further from the center line) in senescent cells, which are not seen or are much reduced in the cycling cells. These changes in interactions appear to be in large domains similar to the size of chromosome bands. Cytogenetic chromosome bands (at 850-band resolution) are defined by the intensity of their staining with Giemsa and are divided into five classes (Gpos100, 75, 50, and 25, and Gneg), which are represented on chromosome ideograms (Fig 7), with “Gpos100” the darkest staining “G-bands” (black), and “Gneg” R-bands (white). We used this chromosome band information (from the UCSC Genome Browser) to map Hi-C interactions, below.

**Figure 7:**
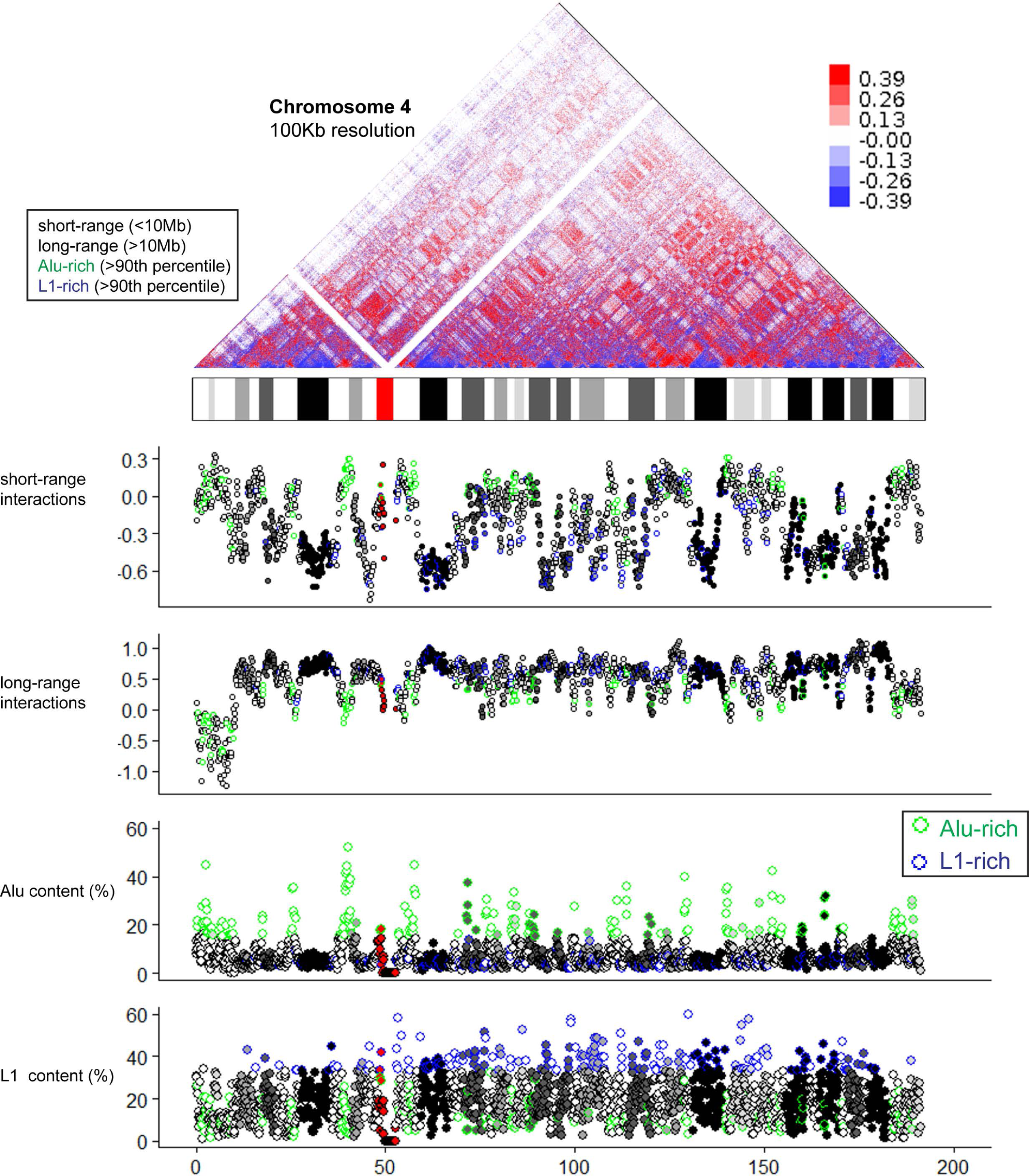
Hi-C Chromosome 4 heatmap of senescent versus growing interactions at 100kb resolution, with ideogram of Chr-4 banding pattern aligned with heatmap below. Below and also aligned to ideogram, is short range (<10Mb) and long range (>10Mb) interactions (Log2 (senescence/growing), and percent Alu content and L1 content. 100kb dots are colored by the 5 Giemsa-band designations and centromere (red), and the dots with highest (90^th^percentile) Alu and L1 are outlined in green and blue respectively.

Chr-4 interaction information between senescent and growing cells is summarized in Figure 7 and 8A, which align all short-(<10Mb) and long-(>10Mb) range interactions with the chromosome ideogram banding pattern, and also with the Alu and L1 density for each contiguous 100 kb of genomic DNA. The 100kb circles on the graph are colored according to chromosome band designation (gray scale) and the circles with highest levels (>90% percentile) of Alu or L1 density are outlined in green or blue respectively. As evident from Figure 7 and 8A, changes in Hi-C interactions occur in large contiguous blocks of genomic DNA, such that numerous adjacent 100kb bins (circles on graph) change in concert, and these mostly correspond to the characteristic cytological-scale banding of Chr-4 seen in the ideogram above. For example, Figure 8A shows a higher magnification view of the short arm of Chr-4, where a large dark band of ∼10 Mb changes as a contiguous unit, with a clear gain in long-range intrachromosomal interactions, which would lessen relative frequency of short-range interactions.

**Figure 8:**
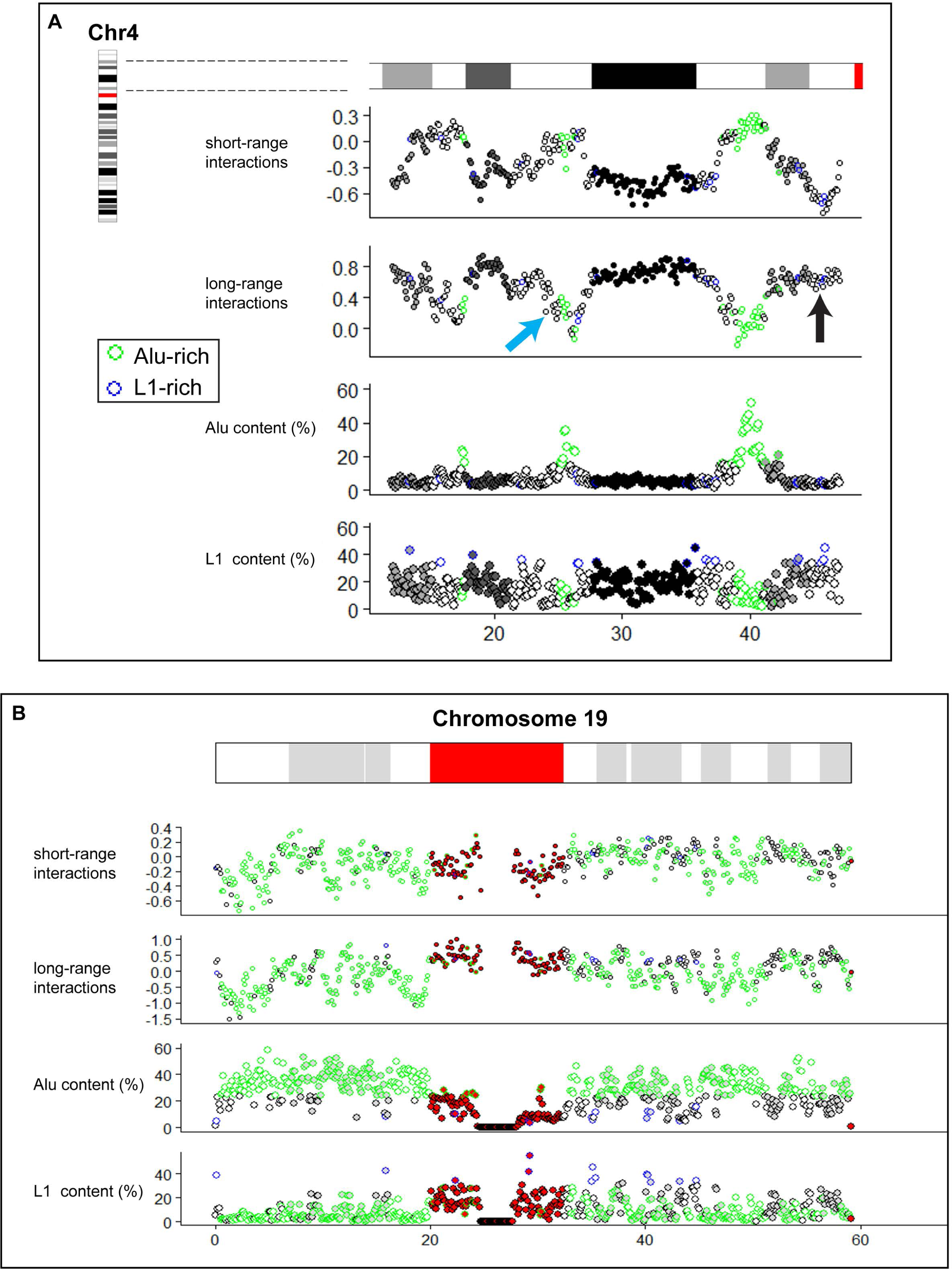
**A**) Close-up of interactions in the short-arm of Chr-4 (not including telomere). Below and aligned to 4p ideogram, is short range (<10Mb) and long range (>10Mb) interactions (Log2 (senescence/growing), and percent Alu content and L1 content. 100kb dots are colored by the Giemsa-band designations, and the dots with highest (90^th^percentile) Alu and L1 outlined in green and blue respectively. Black arrow indicates R-band lacking Alu enrichment. Blue arrow indicates R-band with partial Alu enrichment. **B**) Hi-C interactions for Chromosome 19, aligned with Chr-19 ideogram, starting with short range (<10Mb) and long range (>10Mb) interactions (Log2 (senescence/growing), then percent Alu content and L1 content. 100kb dots are colored by their Giemsa-band designations and centromere (red), and the dots with highest (90^th^percentile) Alu and L1 outlined in green and blue respectively.

A similar general pattern is also evident across all of Chr-4 in Figure 7. The heatmap shows alternating large regions of the chromosome that lose short-range interactions (blue pyramids on the horizontal) and gain long-range interactions (red regions above the horizontal). These large regions correspond to the dark staining G-bands on the Chr-4 ideogram, suggesting that distant syntenic G-bands (known to be L1-rich) are compacting together into SAHFs. If so, one would expect the increase in long-distance interactions seen in G-band DNA to be *with other syntenic G-bands*. This is apparent in the Chr-4 heatmap, where the long-range interactions (red boxes higher in the heatmap) connect all the G-bands along the chromosome (tracing down the left/right diagonals from the boxes). This rearrangement is strongest for the darkest G-bands, which gain the most intra-chromosomal interactions with each other and then to a lesser degree with the lighter G-bands, suggesting the darkest G-bands make up the central core of the SAHF.

Interestingly, the L1 density for Chr-4 is high across the entire chromosome, including both light and dark bands, which do not show sharp differences in L1 content. However, we can still see that the dark G-bands (black-gray dots) lose short-range and gain long-range syntenic interactions in senescent cells, while parts of Gneg R-bands (white dots) do not. Most strikingly, the highest density of Alu DNA on Chr-4 is seen in contiguous stretches that form sharp peaks that reflect marked increases in Alu content over the baseline levels of Alu through most of the chromosome’s dark and light bands. These peaks in Alu density reside in R-bands that are resisting the interaction changes seen on the rest of the chromosome, remaining near zero for both short-range and long-range interactions. (Note: we find R-bands near telomeres behaved differently than most R-bands). These results fit with our DNA FISH data (above) which indicates Alu-rich DNA is depleted from the condensed core of SAHFs and remains more decondensed, to spread across the enlarged nucleoplasm of senescent cells. The resistance of Alu-rich peaks in light R-bands to SAHF coalescence is more apparent in Fig 8A (green outlined white dots), which also suggests that Alu peaks may impact the structural interactions of neighboring sequences. For example, R-bands that lack a “resistant” Alu-peak (Fig 8A, black arrow) can gain long-range interactions similar to darker bands, and when only a portion of white R-bands contain an Alu peak (Fig 8A, blue arrow), the Alu-depleted part of the band may show increased long-range interactions similar to an adjacent dark band.

It is noteworthy that most of Chr-4 has uniformly high L1-density across both light and dark bands, yet still the large dark G-band behaves as one contiguous region (Fig 8A). This suggests to us that some additional property of dark band DNA, other than just L1-density, helps drive or allow for heterochromatin compartment formation. While some level of L1 density may be a pre-requisite, results suggest Alu density, and perhaps relative L1/Alu density, is important. Analysis of Chr-4 indicates that dense Alu peaks resists structural changes that increase long-range intrachromosomal interactions, in contrast to L1-rich/Alu-poor regions, consistent with the latter forming the SAHF. This would predict that for Chr-19, which is remarkably Alu-rich and L1-poor and does not form a SAHF (Fig 5D-F), these extensive changes in long versus short-range interactions would not occur in senescent cells. As shown in Figure 8B, and in sharp contrast to Chr-4, Chr-19 doesn’t show the marked changes in chromosomal interactions; it appears that almost the whole chromosome essentially resists the dramatic reorganization of DNA that is widespread in senescence. All of these Hi-C findings match our *in-situ* observations for Chr-4 and Chr-19 (Fig 5).

The resistance of the Alu-rich regions to SAHF formation remains apparent when examining the combined intrachromosomal short-range and long-range interactions for *all human chromosomes* (at 100kb resolution)(Fig 9). We find essentially all dark band DNA gains long-range syntenic interactions genome-wide, and L1 DNA behaves similarly, consistent with most chromosomes forming L1-rich SAHFs. However, we do see some light R-bands gaining long-range interactions, likely reflecting findings above suggesting R-band DNA lacking Alu-rich peaks could gain long-range interactions because it lacks “Alu-resistance” (Fig 8A; black arrow). This is further supported by findings that Alu-rich DNA (across all human chromosomes) shows little change in long-range intrachromosomal interactions (Fig 9), suggesting the highest Alu-rich regions of the genome resist these changes *en mass*.

**Figure 9:**
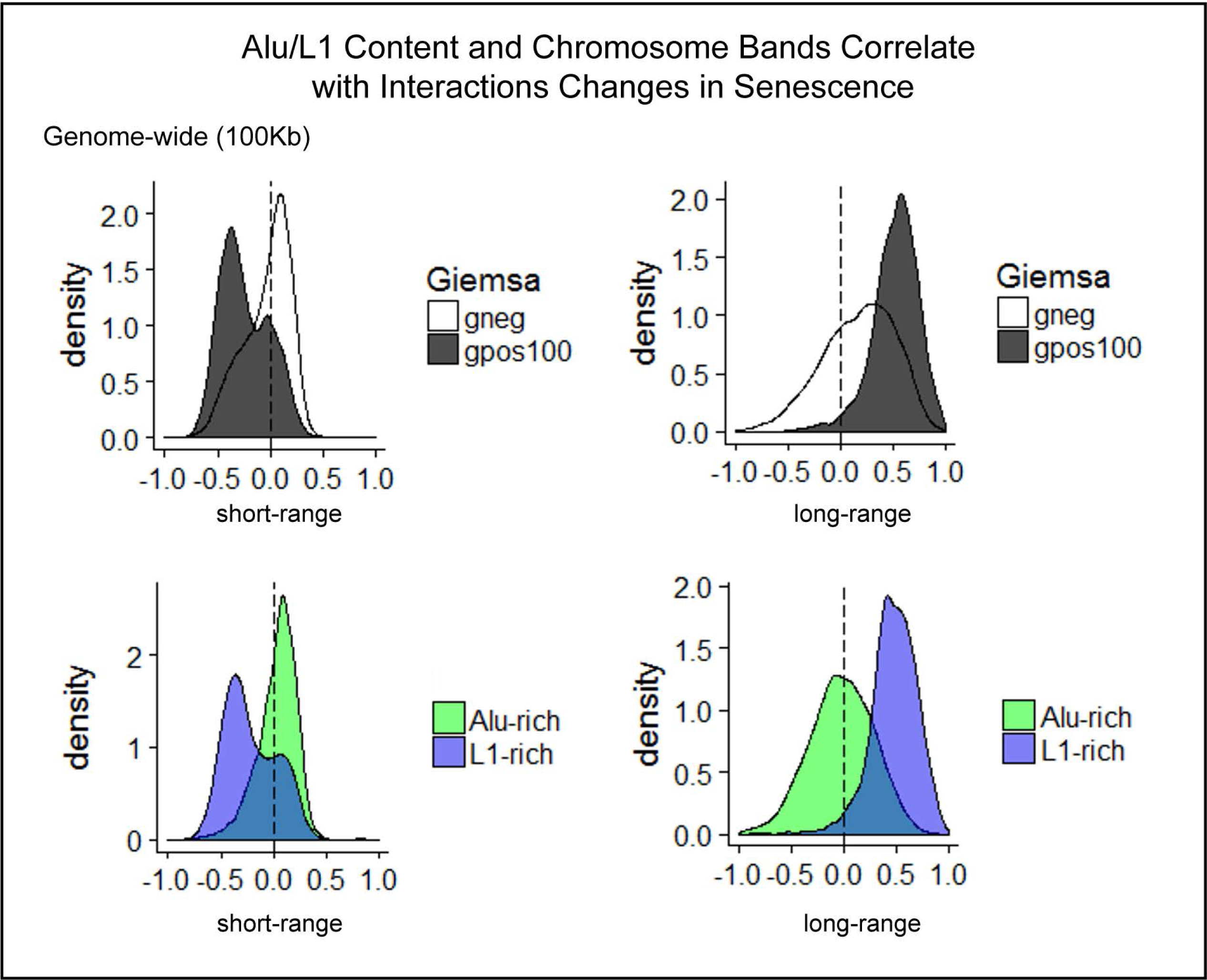
Intrachromosomal Hi-C short-range (<10Mb) and long-range (>10Mb) interactions for all human chromosomes at 100kb resolution for darkest G-bands (Gpos100) and light R-bands (Gneg) on top, or highest (90^th^ percentile) Alu (green) or L1 (blue) DNA below.

In sum, results show L1-rich dark band DNA gains long-range interactions, and the highest Alu-rich peaks within light bands really resist this change. This suggests that Alu-rich DNA possesses a property that keeps it from coalescing into the dense, silent heterochromatin compartments that can form in the chromosome territory (BB or SAHFs). The fact that Alu-rich peaks in the uniformly L1-dense Chr-4 can resist the dramatic changes that form SAHFs, may be facilitated by association of active genes with Nuclear Speckles (reviewed in: Smith et al., 2020), as we observe Alu-rich DNA is still often closely associated with SC35 domains in the enlarged nucleus of senescent cells.

## DISCUSSION

This study provides new insights into what we suggest is a fundamental relationship between the segmental organization of genomic sequences along chromosomes and interphase nuclear genome organization, that is linked to gene regulation. Metaphase chromosomes exhibit clear organization of sequence content that is reflected in dark G-band and light R-band staining patterns, any functional implications of which are currently unknown. Perhaps because of the invariant and very large-scale nature of this pattern, these cytologically distinct blocks of linear DNA have received little to no consideration in most current studies of nuclear genome architecture. Our genome also contains a vast amount of repetitive DNA of different types that show distribution differences that largely correlate with chromosome bands, and similar syntenic blocks of DNA are largely conserved across species. Here we utilize two model systems that allow us to examine changes in these distinct repeat-rich regions across a single chromosome territory, with XIST RNA mediated chromosome silencing (Barr body formation of the Xi), and across an entire genome, with formation of senescent associated heterochromatin foci (SAHFs) in senescent cells. Results show that the two most abundant repeats in our genome, L1 and Alu, tend to segregate within chromosome territories, and between large nuclear compartments in interphase nuclei. While on a smaller scale, this “homotypic clustering” of repeats could largely reflect the linear organization of Alu and L1 sequences that pre-exist on chromosomes. However, this study goes further to show that these large regions (bands) of DNA with similar L1/Alu content self-associate *across the whole chromosome territory* in these model systems, producing essentially two large domains of coalesced syntenic L1-rich/Alu-poor chromatin and Alu-rich chromatin. In both these model systems, this coalescence of large L1-rich blocks of DNA forms a silent (CoT-1 RNA depleted) domain at the core of each chromosome territory surrounded by gene-rich DNA.

Importantly, results here show for the first time that the DNA that forms this heterochromatic core is not only L1 dense, but, most notably, is sharply depleted of Alu-rich regions. Contiguous stretches of DNA that are the most highly enriched for Alu were seen to withstand the apparent widespread “collapse” of L1-rich chromatin into the visibly condensed SAHF structures. This resistance of the most Alu-rich regions to whole chromosome condensation is likely linked to its relationship with highly expressed genes that associate with euchromatic nuclear structures, most notably nuclear speckles (aka: SC-35 domains). Even for the Xi in which most genes are silenced, the Alu-rich DNA remains largely peripheral and at the periphery of the L1-rich chromosome core, the condensed Barr Body. These findings have relevance to both XIST RNA function during X-chromosome inactivation as well as gene regulation globally.

### Formation of BB/SAHFs is not simply due to silencing of canonical genes and their condensation at the histone-nucleosome level

The Barr body (BB) of the inactive X-chromosome (Xi) (and to some extent the SAHFs in senescent cells) were long presumed to be the visible manifestation of silenced genes, compacted by accumulated epigenetic (histone/nucleosome) modifications to produce the condensed structure seen by DNA stain (Chaligne and Heard, 2014; Lee and Bartolomei, 2013; Narita et al., 2003). However, canonical gene silencing would only impact a minor fraction of chromosomal DNA, hence we have long argued that the DNA-dense BB is huge relative to what would be expected from silencing the subset of X-linked genes active in any given cell type (Hall and Lawrence, 2010). In fact, reports suggest there is little difference in the amount of “open chromatin” between the active and inactive X-chromosome (Xa & Xi) (Gilbert et al., 2004; Naughton et al., 2010; reviewed in: Pandya-Jones and Plath, 2016), which was limited to the TSS of silenced genes and thus could not be responsible for the large-scale condensation seen for the BB. In addition, we have previously reported that XIST RNA starts to build the condensed BB structure *days* before it silences canonical genes (Valledor et al., 2023). Our findings here fit well with this timing difference, since the BB structure is composed of gene-poor, L1-rich DNA rather than gene-rich Alu-rich DNA. These results collectively suggest that XIST RNA acts to compact L1-rich/Alu-poor and gene-poor DNA first, concentrating it in a dense silent core (BB) that is formed before most canonical genes in Alu-rich regions become silenced. This further explains why canonical genes in Alu-rich DNA are localized at the periphery of the dense BB regardless of silenced state (Clemson et al., 2006). We also find that XIST RNA binds *both* Xi domains, that contain different repeat composition (L1/Alu), gene densities (gene-rich/gene-poor), and epigenetic histone marks (H3K27me3/H3K9me3), which is contrary to some publications (e.g. Chadwick and Willard, 2004; Nozawa et al., 2013; reviewed in: Dixon-McDougall and Brown, 2016). However, we find that whether XIST RNA paints H3K9me3 labeled DNA, L1-rich DNA, or G-bands may ultimately be affected by the state of the cells (mitotic, cancer, senescence, transcriptional), choice of antibody and probe and/or labeling conditions, rather than the native binding of XIST RNA.

Senescent cells also form large condensed L1-rich/Alu-poor, gene-poor cores at the center of most chromosomes and they similarly lack CoT-1 RNA and Alu-rich DNA. Although cell-cycle genes are specifically silenced in senescent cells, interestingly, expression changes in senescent cells are mostly associated with *new* expression of many genes (e.g. SASP, cytokines, chemokines, and growth factors) rather than genome wide gene silencing (Hernandez-Segura et al., 2017; Marthandan et al., 2016). Consistent with this, in senescent cells there is not only SAHF condensation, but an expansion of highly *decondensed* chromatin between SAHFs in a much-enlarged nucleus, including a marked distension of normally compact centromeric satellite DNA (Swanson et al., 2013). This and other findings have indicated that changes to genome packaging in senescent cells also occurs at a level above nucleosome/histone modifications on silenced genes; for example, it has been reported that SAHF formation occurs without major changes at the local level to repressive marks (H3K9me3 and HeK27me3) between growing and senescent cells (Chandra et al., 2012).

### L1-rich chromatin condensation may be facilitated by lack of euchromatin-associated RNAs and loss of scaffold attachment

Both SAHFs and BBs lack euchromatin associated RNAs (e.g. CoT-1 RNA), which are thought to promote open chromatin structure through RNA-dependent links to the nuclear scaffold (Creamer et al., 2021; Hall et al., 2014). It’s hypothesized that removal of these often lowly expressed, repeat rich RNAs alters chromatin association with the nuclear scaffold and results in DNA condensation on a histological scale (rather than at the histone/nucleosome level). It is possible that the SAHF/BB structure could be formed from already silent L1-rich DNA, making the Cot-1 RNA/RNAPII “hole” over these structures a consequence of L1 DNA re-organization. However, another possibility (not mutually exclusive) is that XIST RNA silences low-level expression of lncRNAs in L1-rich/gene-poor DNA which causes it to collapse into a large condensed nuclear structure.

For example, the action of XIST RNA on L1-rich DNA early in the inactivation process may function largely through loss of lncRNAs and other low-level repeat-rich transcripts in the RNP scaffold, disruption of which has been shown to cause cytological-scale DNA compaction (Creamer et al., 2021). This is supported by other reports that find XIST RNA adds repressive histone marks (H3K27me3 & H2AK119ub) on gene-poor “*intergenic”* DNA first, before moving onto more active “genic” regions (Zylicz et al., 2019). Loss of low-level transcripts, that impact the RNP scaffold at cytological scale, would also fit with our finding here that the condensed BB often exhibits reduced staining for Lamin B1, a nuclear scaffold protein. Further suggestive of nuclear scaffold changes being associated with large-scale structural changes, loss of Lamin B1 throughout the nucleus is also a characteristic of senescent cells (Sadaie et al., 2013; reviewed in: Lukasova et al., 2018). Thus, XIST RNA may mediate changes in Lamin B1/L1-rich DNA interactions for a *single chromosome* that is paralleled, genome-wide, in senescent cells.

### An L1-rich gene-poor compartment will favor sustained histone de-acetylation required for subsequent steps in stable epigenetic silencing

In numerous earlier papers we showed that many highly active genes were not only in the euchromatic nuclear compartment, but specifically localized at the periphery of an SC-35 rich nuclear speckle (reviewed in: Hall et al., 2006; Smith et al., 2020). It has since become widely accepted that localization of active genes with domains of concentrated pre-mRNA metabolic factors will facilitate gene expression. However, why form a “compartment” for silenced genes since epigenetic regulation of promoters is known to provide stable silencing? Insight into this came from a recent study (Valledor et al., 2023) in which we showed that a required step to initiate gene silencing, H3K27 de-acetylation, is a highly *unstable* step that can be rapidly reversed by histone acetyltransferases (HATs), which are prevalent in active chromatin regions. Results showed that high cytological density of XIST RNA (or small ncRNAs containing the silencing A-repeat domain) was necessary to shift the ongoing balance of HATs and HDACs (Histone deacetylases) toward deacetylation, allowing subsequent epigenetic steps to stabilize the silent state. We propose that formation of the large Barr Body, enriched for transcriptionally silent, gene-poor, L1-rich DNA will lack substantial HAT activity and thus provide a “silencing compartment” which favors effective/persistent histone deacetylation. Effective H3K27-deacetylation will then allow methylation of the same residue (PRC2 mediated H3K27me3), and subsequent steps, which stabilize the silent state. Thus, although SAHF-containing chromosomes in senescent cells have much more gene expression than the inactive X-chromosome, we propose that proximity of canonical gene promoters to the edge of the HAT-depleted BB/SAHF structure facilitates sustained gene silencing (cell cycle genes or silenced X-linked genes) by preventing re-acetylation of genes that had been in active chromatin regions, facilitating stabilization of the silenced state (illustrated in Figure 10). In contrast, regions of active, gene-dense, Alu-rich DNA in HAT enriched euchromatic regions, particularly regions associated with nuclear speckles, ensure stable acetylation and activity of highly active genes (or escaping X-linked genes). This might suggest that genes enriched for L1 DNA might exhibit enhanced silencing due to proximity to the silencing domain, while genes highly enriched for Alu DNA might resist tight association with the silencing domain and partially or fully escape silencing. This is supported by reports from X:autosome translocation studies that find genes effectively silenced by XIST RNA are enriched for L1-sequences, whereas those that escape silencing contain significantly more Alu-sequences (Cotton et al., 2014).

**Figure 10:**
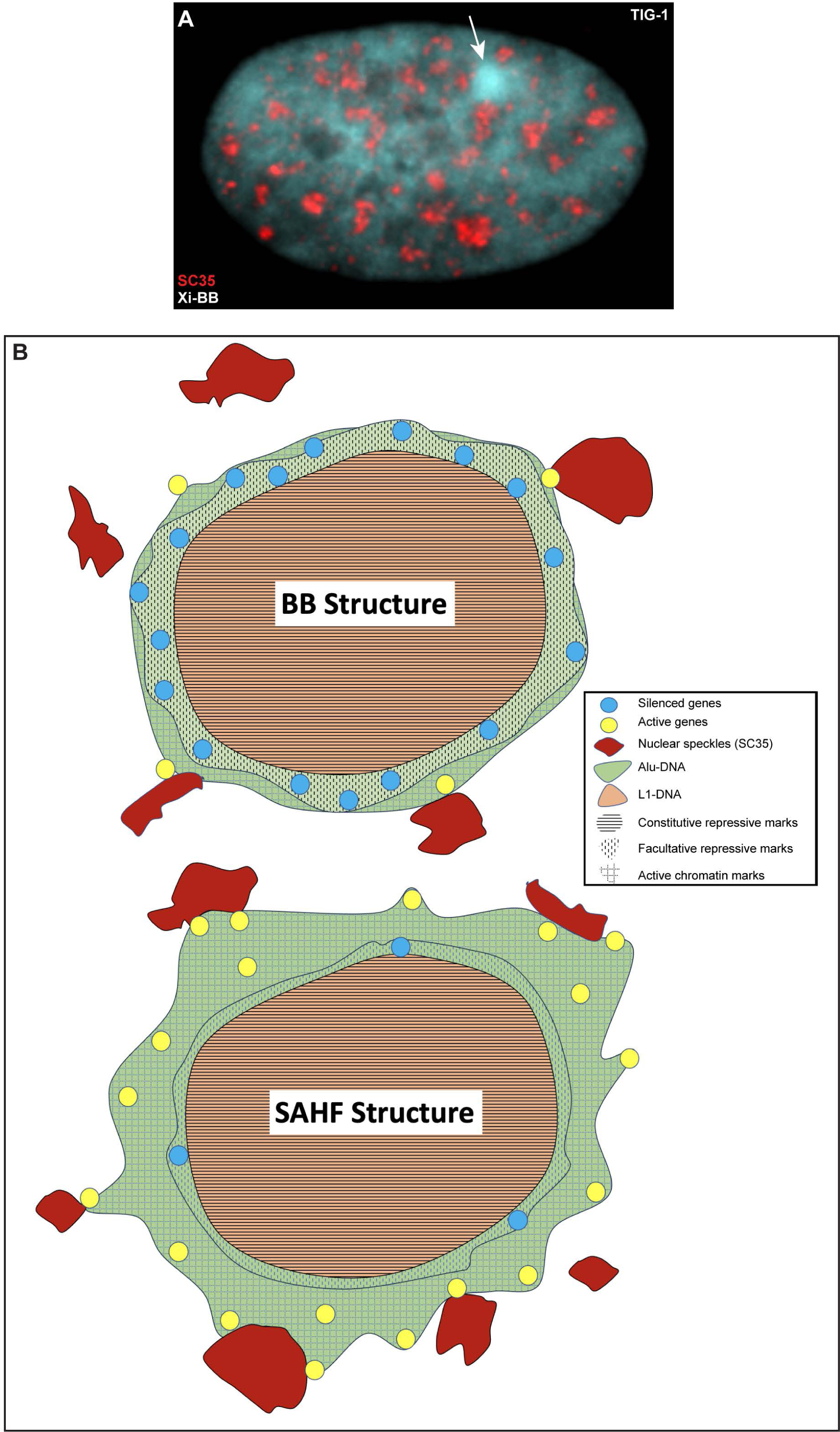
**A**) Growing female fibroblast with DAPI (white) and SC35 protein (nuclear speckles) red, showing the BB (arrow) surrounded by nuclear speckles. **B-C**) Illustration of BB (**B**) and SAHF (**C**) chromatin structure.

### A hypothesis: Segmental DNA organization on chromosomes facilitates regulated genome function within compartmentalized nuclear structure

This study provides evidence that very large segments of DNA, corresponding loosely to dark versus light chromosome bands, undergo packaging changes as a collective unit. During SAHF formation in senescent cells (and likely BB formation of the Xi), gene-poor dark-band DNA collapses into the central core of the chromosome territory to coalesce with other syntenic dark-bands to form a heterochromatic SAHF. Importantly, these condensed gene-poor cores are depleted of Alu-rich DNA. In fact, the most Alu-rich segments of the chromosome resist this structural change *as a unit.* Similarly, large (∼5-10 Mb) blocks of dark-band DNA undergo changes in long-range interactions as a contiguous unit as well. This suggests large contiguous chromosomal domains contain similar sequence properties that facilitate either coalescence or resistance to cytological condensation.

A question remains whether the L1 or Alu repeats themselves play active or passive roles in this segregation, as their genomic distribution does not reflect a preference for insertion site, this suggests some evolutionary selection for genomic distributions in chromosome bands (Sultana et al., 2019). L1 sequences are thought to contain features that form unusual DNA structures (Usdin and Furano, 1989), which can be modulated by changes in PH, cations, histones, density of repeats, and torsional stress, promoting self-association. Whereas SINES are shown to act as boundary-elements/buffers/insulators (reviewed in: Lunyak and Atallah, 2011), and we see Alu-rich peaks anchoring the gene-rich chromosome bands that resist compaction into SAHFs. Thus, they may buffer/insulate the “spread” of heterochromatin by physically separating active DNA from silent L1-rich DNA within nuclear/chromosome structure. However, we also find that many chromosomes contain fairly uniform L1-density across all bands, yet large G-bands still behave as one unit compared to neighboring lighter bands. Thus, although there may be some level of L1-density required, it’s possible that another sequence property of dark-band DNA may actually drive coalescence of the whole contiguous region. Similarly, Alu-rich DNA resistance to coalescence may be more due to gene activity and the presence of euchromatic-associated RNAs than anything inherent to the Alu-sequence itself. It therefore remains to be established whether L1 and Alu repeats themselves determine this regional unity or other aspects of the sequence content or function play a role.

Finally, we propose that chromosomal bands, which are enriched for different repeats and gene densities, are a preexisting structural framework for the *regional* (mega-base scale) regulation of the genome, and the nuclear architecture it builds. This constitutive framework may be “hiding in plain sight”, in the form of very large blocks of differently staining DNA on mitotic chromosomes that we have spent the last 60 years observing but not understanding (or even studying).

## ACKNOWLEDGMENTS

We appreciate the support of the NIH-grants R01R35GM122597, R01HD091357, and R01HD094788 to J.B.L. We thank members of the Lawrence lab for support in various ways, including thoughtful discussions and critical analysis of this study, and our summer interns Alana Anderson and Maya Kasbekar for performing the X-linked gene DNA, XIST RNA FISH experiments.

## METHODS

### Cell lines, growth conditions & fixation

All cell lines (List in Supplement), were grown in conditions recommended by suppliers (See supplemental Methods). Drug treatments (STSP & DRB) are detailed in supplement. Our standard fixation protocols have been detailed previously (Byron et al., 2013; Johnson et al., 1991; Tam et al., 2004) (details in Supplement).

### FISH and IF

Our standard hybridization conditions for RNA, DNA, simultaneous DNA/RNA, and simultaneous DNA or RNA and IF detection was performed as previously described (e.g. (Byron et al., 2013)) (details in Supplement). Specific probes, chromosome paints and antibodies are listed in Supplement. Nick-translated genomic probes were used for most images unless otherwise indicated.

### Microscopy and Digital Imaging

An Axiovert 200 or an AxioObserver 7 Zeiss microscope equipped with a 100X PlanApo objective (NA 1.4) and Chroma 83000 multi-bandpass dichroic and emission filter sets (Brattleboro, VT. Images and Z-stacks were captured with the Zeiss AxioVision or Zen software, and an Orca-ER camera or with a Flash 4.0 LT CMOS camera (Hamamatsu). Images were minimally corrected for contrast and brightness (min/max), to best represent signals observed by eye using Axiovision or Zen (Zeiss) software, unless otherwise noted in figure legend or text. Where required, care was taken to eliminate any bleed-thru of red fluorescence into the fluorescein channel. Images are 2D, show a plane from the z-stack or a MIP (as indicated). Most experiments were carried out a minimum of 2 times, with typically 100-300 cells scored in each experiment. Key results were confirmed by at least two independent investigators. All findings were easily visible by eye through the microscope (unless otherwise noted), and images were minimally enhanced for brightness and contrast in Photoshop to resemble what was seen by eye through the microscope (unless otherwise noted).

### L1 Alu content in Chromosomes

Contribution of L1 and Alu sequences to human chromosomes were calculated for the hg19 Feb 2009 human genome reference assembly (hg19) using the RepeatMasker repeat library (v4.0.5).

### Hi-C analysis

Published Hi-C datasets were downloaded from accession PRJEB80 (PMID 25640177). Reads were mapped using Bowtie to the human genome (hg19) and contact matrices were generated using Homer at the indicated resolution (PMID 20513432). Options -*removePEbg -restrictionSite AAGCTT -removeSelfLigation - removeSpikes 10000 5 parameters were employed when using the makeTagDirectory* tool. Percent LINE-1 and Alu sequences in each genomic interval of the count matrices was calculated using Repbase annotations (downloaded from the UCSC browser) and BEDtools. Chromosome band information was downloaded using the UCSC Genome Browser.

