## Supplemental Figures and Methods for "Differences in Alu vs L1-rich chromosome bands underpin architectural reorganization of the inactive-X chromosome and SAHFs"

Supplemental Figure 1

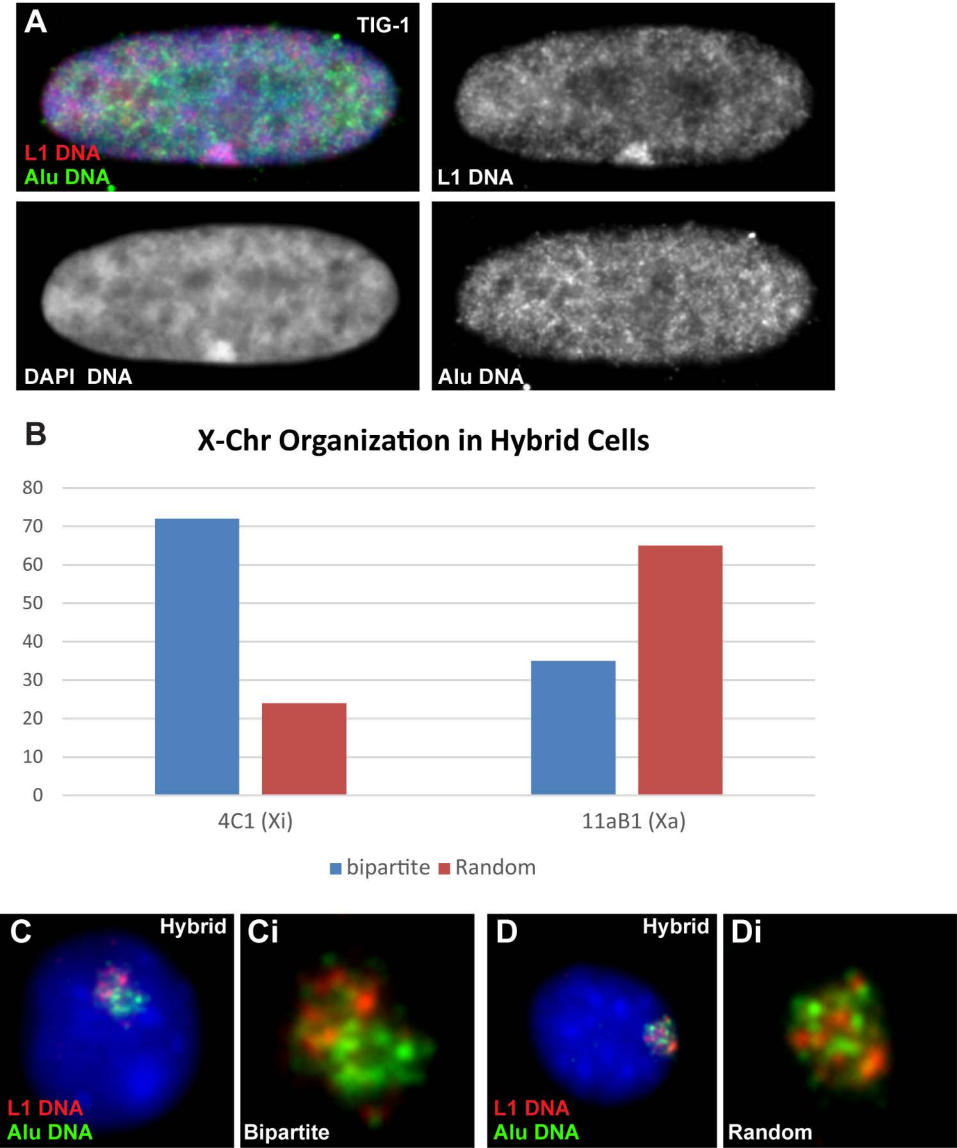

Supplemental Figure 2

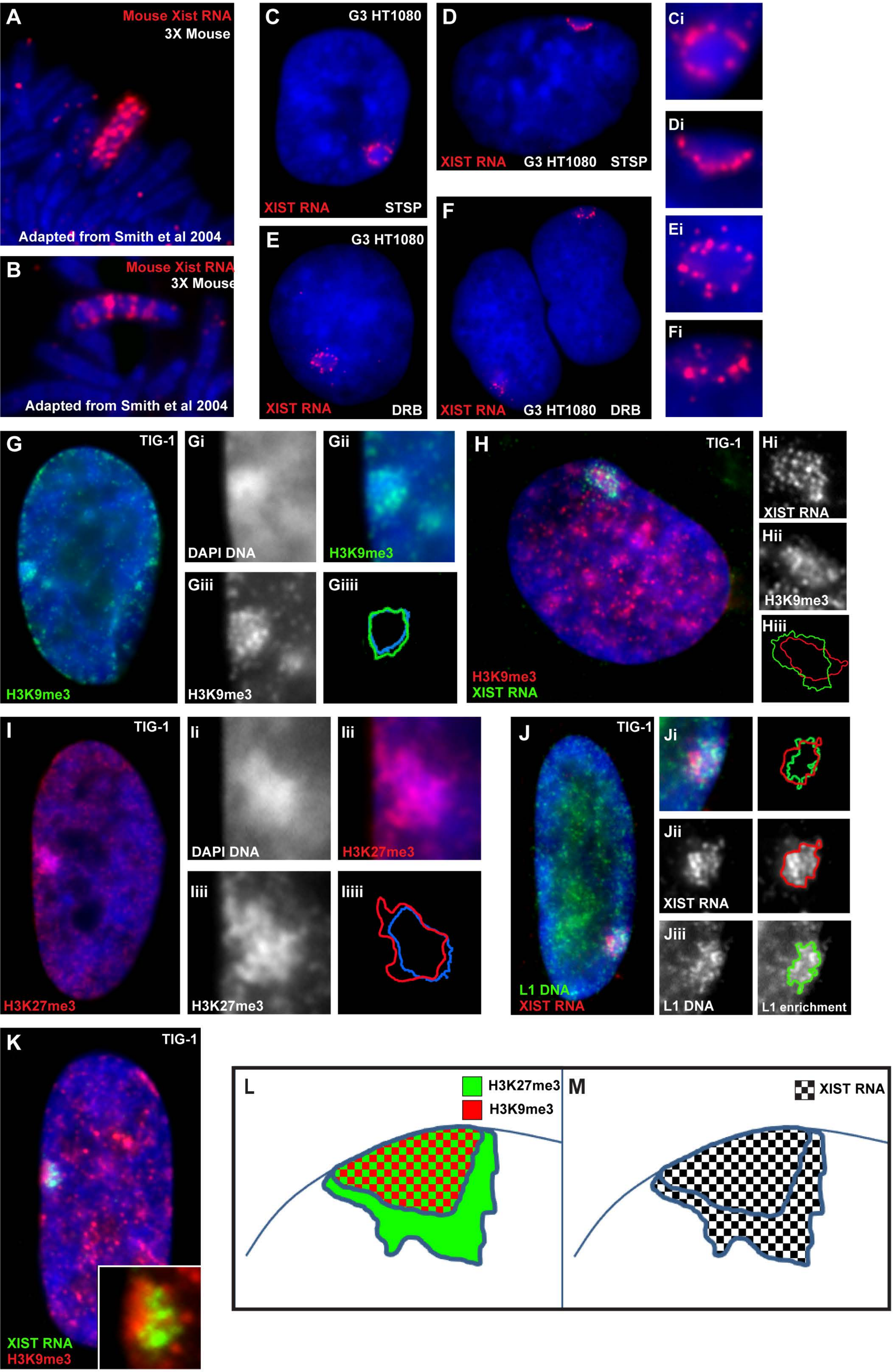

Supplemental Figure 3

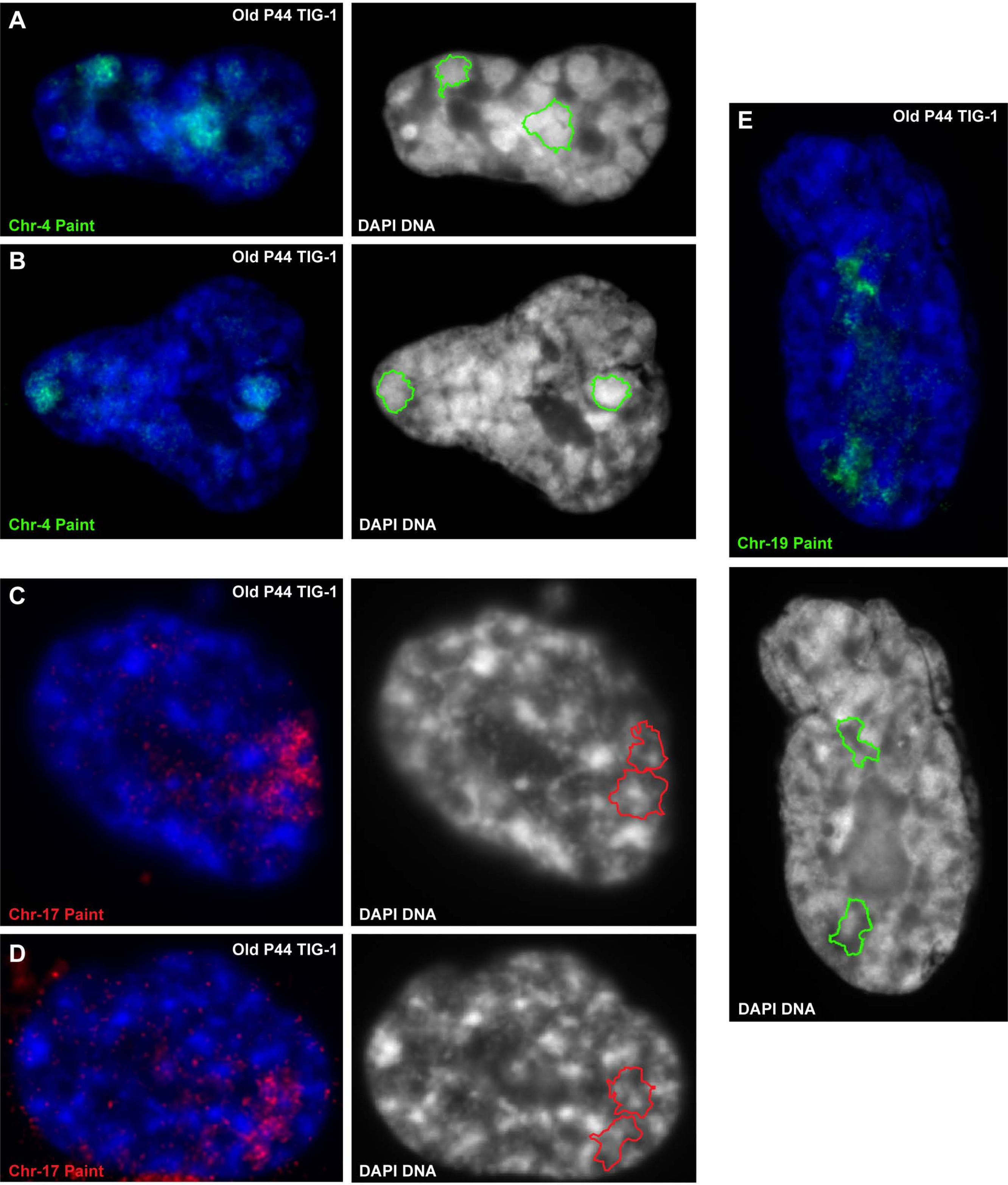

Supplemental Figure 4

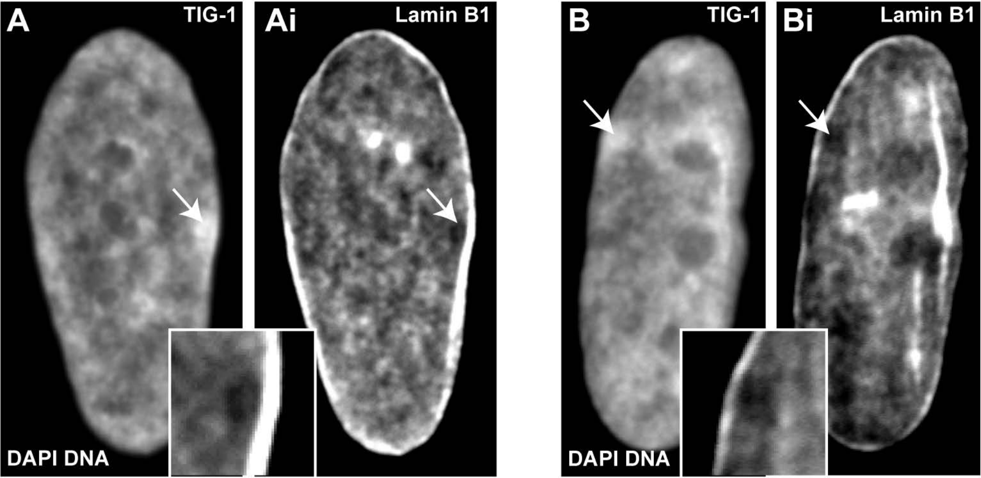

### SUPPLEMENTAL FIGURE LEGENDS

**Supplemental Figure 1**: DAPI DNA stain (blue) used in all images. **A**) Female fibroblast with L1 DNA (red) and Alu DNA (green) FISH. Separated channels included. **B-D**) Mouse/human hybrid cell lines with a single human inactive X-chromosome (Xi) (4C1) or a single active X-chromosome (Xa) (11aB1) were scored for L1 or Alu organization, either in two-domains (bipartite) or intermixed (random) (n=100 for each of 2 experiments). Images of representative “bipartite” or “random” cells are shown below (**C-D**). Two-color enlarges of human chromosome at right.

**Supplemental Figure 2**: DAPI DNA stain (blue) used in all images. **A-B**) Mouse mitotic chromosomes with mouse Xist RNA FISH, showing complete coverage of the mouse Xi (**A**) or residual coverage on light R-bands (**B**). Images were adapted from those we used in (Smith et al., 2004). **C-F**) Human G3 HT1080 cells treated with STSP (**C-D**) or DRB (**E-F**) for 5 hours exhibit residual XIST RNA (red) on DNA around the outer edge of the BB. Enlargement of the BB regions in each cell at right. **G**) H3K9me3 antibody (Upstate: 07-422) labels the BB of the Xi in female fibroblasts. Enlargement and separated channels of the BB region at right. The areas of DAPI intensity and H3K9me3 enrichment are outlined and overlaid with each other to show overlap. **H**) Using the full-length (15kb) cDNA XIST probe for RNA FISH, XIST RNA (green) labels the Xi in female fibroblasts including DNA enriched for H3K9me3 (red). Enlarged and separated channels of the BB region at right, with areas of XIST RNA and H3K9me3 enrichment outlined and overlaid to show overlap. **I**) H3K27me3 antibody (Millipore: 07-449) is usually slightly larger than the DAPI dense BB of the Xi in female fibroblasts. Enlargement and separated channels of the BB region at right, with areas of DAPI intensity and H3K27me3 enrichment outlined and overlaid to show overlap. **J**) XIST RNA (red) paints the L1-rich (green) BB of the Xi in female fibroblasts. Enlargement and separated channels of the BB region at right, with areas of XIST RNA and L1 DNA enrichment outlined and overlaid to show overlap. **K**) Using the small (9kb) 3’-genomic G1A XIST probe (available at Addgene) for RNA FISH, XIST RNA (green) does not paint H3K9me3 enrichment chromatin (red) in female fibroblasts, suggesting G1A probe hybridization is blocked (masked) in H3K9me3 labeled DNA. Two-color closeup of BB region (insert). **L**) Diagram illustrating the Xi with L1-rich BB containing both H3K9me3 and H3K27me3, while Alu-rich exterior labels with H3K27me3, and XIST RNA paints both Xi domains.

**Supplemental Figure 3:** DAPI DNA stain (blue) used in all images. **A-B**) Senescent fibroblast with Chr-4 DNA FISH (green). Separated DAPI channels at right with outlines of both Chr4 territories superimposed on DAPI image. **C-D**) Senescent fibroblast with Chr-17 DNA FISH (red). Separated DAPI channels at right with outlines of both Chr-17 territories superimposed. **E**) Senescent fibroblast with Chr-19 DNA FISH (green). Separated DAPI channel below with outlines of both Chr-19 territories superimposed.

**Supplemental Figure 4: A-B)** Separated channels of DAPI DNA stain (left) and Lamin B1 staining (right) in growing female fibroblasts, with BBs indicated by arrows. Inserts are an enlargement of region over BB, from Lamin B1 channel.

### SUPPLEMENTAL METHODS

**Cell culture and treatments**: Young TIG-1 human fibroblasts (Coriell Institute for Medical Research, NIA Aging Cell Culture Repository) were cultured in Minimal Essential Media (MEM) supplemented with 15% FBS and passaged 1:2 as cultures reached confluency. Senescent TIG-1 fibroblasts were defined as senescent when they failed to reach confluency 10 d after a 1:2 split, usually around passage 44-45, and could be maintained in this state for several weeks. Neoplastic transgenic line, G3-HT1080 (Hall et al, 2002), was grown in DMEM with 10% FBS. Mouse-Human hybrid cells, 4c1 (Xi) and 11aB1 (Xa), (Gift from Carolyn Brown, University of British Colombia) were grown in Minimal Essential Media (MEM) supplemented with 10% Fetal Bovine Serum. Human H9 ES cells (WiCell) were grown on irradiated mouse embryonic fibroblasts (iMEFs) (R & D Systems, PSC001) in hESC medium containing DMEM/F12 supplemented with 20% knockout Serum Replacement (Invitrogen), 1mM glutamine (Invitrogen), 100 μM non-essential amino acids (Invitrogen), 100 μM β-mercaptoethanol (Sigma) and 10 ng/ml FGF-β (Invitrogen, PHG0024). Cultures were passaged every 5–7 days with 1 mg/ml of collagenase type IV (Invitrogen). Transcription inhibitors were used briefly to illustrate some points: 6-dichloro-1-β-D-ribofuranosylbenzimidazole (DRB), was used at 40 μg/ml for 4 hrs and staurosporine (STSP) at 1μM for 5 hours. Drugs were dissolved in DMSO for stock solutions, then added directly to media over monolayer cells on coverslips.

**Cell Fixation:** Cultured cells were grown on glass coverslips prior to fixation. Our standard fixation conditions used in most experiments has been previously published (Byron et al., 2013). Briefly, cells on glass coverslips are extracted in CSK buffer, 5% triton, and VRC (vanadyl ribonucleoside complex) for 1-3 min. Cells were fixed in 4% Paraformaldehyde for 10min, then stored in 1XPBS or 70% ETOH.

**RNA and DNA FISH & IF:** Our standard hybridization conditions for RNA, DNA, simultaneous DNA/RNA, and simultaneous DNA/IF or RNA/IF detection was performed as previously described (Byron et al., 2013), and briefly described below. All slides were counter stained with DAPI. Vectashield (Vector Labs) was used as mounting media for all fluorescence imaging. Larger (non-oligo) DNA probes were nick translated with biotin-11-dUTP or digoxigenin-16-dUTP (Roche Diagnostics, Indianapolis, IN). Hybridizations were overnight at 37°C, in 2xSSC, 1U/ul RNasin and 50% formamide, with 2.5ug/ml of DNA probe. Cells were washed: 15% formamide/2xSSC at 37°C (20min); 2xSSC at 37°C (20min); 1xSSC at RT (20min); and 4xSSC at RT (5min). Detection utilized Alexa 488 or Alexa 549 Streptavidin (Invitrogen) in 1%BSA/4xSSC for 1 hr at 37°C. Post-detection washes: 4xSSC; 4xSSC with 0.1% Triton; and 4xSSC, each for 10 min at RT, in the dark. Unlabeled Human CoT-1 DNA (10-15mg) was included in the hybridization buffer to block non-specific background in all RNA FISH reactions except for CoT-1, L1 or Alu RNA hybridizations. Chromosome paint hybridizations performed following manufacturer's instructions.

Simultaneous RNA/DNA hybridizations: To detect RNA exclusively, all RNA hybridizations were carried out under non-denaturing conditions, preventing DNA accessibility to the probe. RNA hybridization was performed before DNA denaturation and hybridization (as described above). Following RNA hybridization, the cells were fixed (4% Paraformaldehyde) for 10 min, then treated with 0.2N NaOH in 70% ETOH for 5 min, rinsed with 70% ETOH then denatured in 70% formamide, 2xSSC, at 75°C for 2 min, up to 5 min for L1 hybridizations, before ethanol dehydration, and air-drying. Then DNA was hybridized and detected as described above.

Simultaneous RNA/DNA and antibody detection: Most antibodies were used prior to RNA or DNA hybridization. Briefly, slides were incubated in the appropriate dilution of primary antibody in 1%BSA, 1xPBS and 1U/ul RNasin, for 1 hour at 37°C. Slides were washed, and immunodetection was performed using 1:500 dilution of appropriately conjugated (Alexa 488 or Alexa 594, Invitrogen) secondary (anti-goat, mouse or rabbit) antibody, in 1xPBS with 1% BSA. The antibody signal is fixed in 4% paraformaldehyde for 10 min prior to hybridization (performed as detailed above).

Probes used: Full length LINE1 (John Moran, UMich), Alu (pPD39, ATCC), XIST: 10kb G1A (Addgene plasmid #24690) and Full length expression plasmid (Valledor et al 2023), X-linked genes: PGK1, ZFX, F8C, SMC1L1 (Clemson et al 2006), & XACT (BAC RP11-35D3), Cot-1 repeat probe (Roche), Chromosome paints: Chr-4 paint and Chr-19 paint (ID Labs Biotechnology, Ontario).

Antibodies used: SC35 (Sigma, S4045), Lamin B1 (Abcam & Santa Cruz), H3K9me3 (Upstate: 07-422), H3K27me3 (Millipore: 07-449).

**Microscopy and Digital Imaging:** An Axiovert 200 or an AxioObserver 7 Zeiss microscope equipped with a 100X PlanApo objective (NA 1.4) and Chroma 83000 multi-bandpass dichroic and emission filter sets (Brattleboro, VT. Images and Z-stacks were captured with the Zeiss AxioVision or Zen software, and an Orca-ER camera or with a Flash 4.0 LT CMOS camera (Hamamatsu). Images were minimally corrected for contrast and brightness (min/max), to best represent signals observed by eye using Axiovision or Zen (Zeiss) software, unless otherwise noted in figure legend or text. When required, care was taken to eliminate any bleed-thru of red fluorescence into the fluorescein channel. Images are 2D, a single plane from the z-stack or a MIP (as indicated). Most experiments were carried out a minimum of 2 times, with typically 100-300 cells scored in each experiment. Key results were confirmed by at least two independent investigators. All findings were easily visible by eye through the microscope (unless otherwise noted), and images were minimally enhanced for brightness and contrast in Photoshop (min/max) to resemble what was seen by eye through the microscope (unless otherwise noted).
